# Ecotypes matter: photoperiod and site of origin underlie the onset of leaf senescence in *Eriophorum vaginatum*, an Arctic foundation species

**DOI:** 10.64898/2026.11.18.766787

**Authors:** Marcel Caritá Vaz, Wesley Mahler, Bjorn Larson, Luis A. Pallares-Solano, Derek D. Jolley, Brenda Del Cid Ramirez, Hashel Orquiz, Caleb Edwards, Emma Klusmeyer, Jianwu Tang, Gaius Shaver, Michael L. Moody, Ned Fetcher

## Abstract

The rapid warming of the Arctic is causing the growing season to expand and the climate to become more unpredictable. This scenario is intensifying a crucial trade-off arctic plants face: maximizing growth during the short growing season while avoiding freezes at the shoulder seasons. In response, ecotypes have evolved phenological behaviors that are finely tuned to their local climate, which often involves the use of photoperiod cues, especially in the fall. Under the rearrangement of temperature and light regimes, however, such phenological adaptations might prevent native plants from taking advantage of longer growing seasons. To test this phenological mechanism of local adaptation we used a decade-long reciprocal transplant experiment in Northern Alaska and observed the effects of seasonal light and temperature regimes on the leaf phenology of an arctic foundation species, *Eriophorum vaginatum*. We found that ecotypes rely heavily on site-specific photoperiod cues, but may also use temperature cues. These results indicate that local adaptation of this widespread species to local climates required the evolution of diverging phenological responses to seasonal changes in light. Because light cues will remain stable as the shoulder seasons expand, these adaptations are increasingly maladaptive, and could leave *Eriophorum* vulnerable to more competitive species.

**Graphical abstract:** 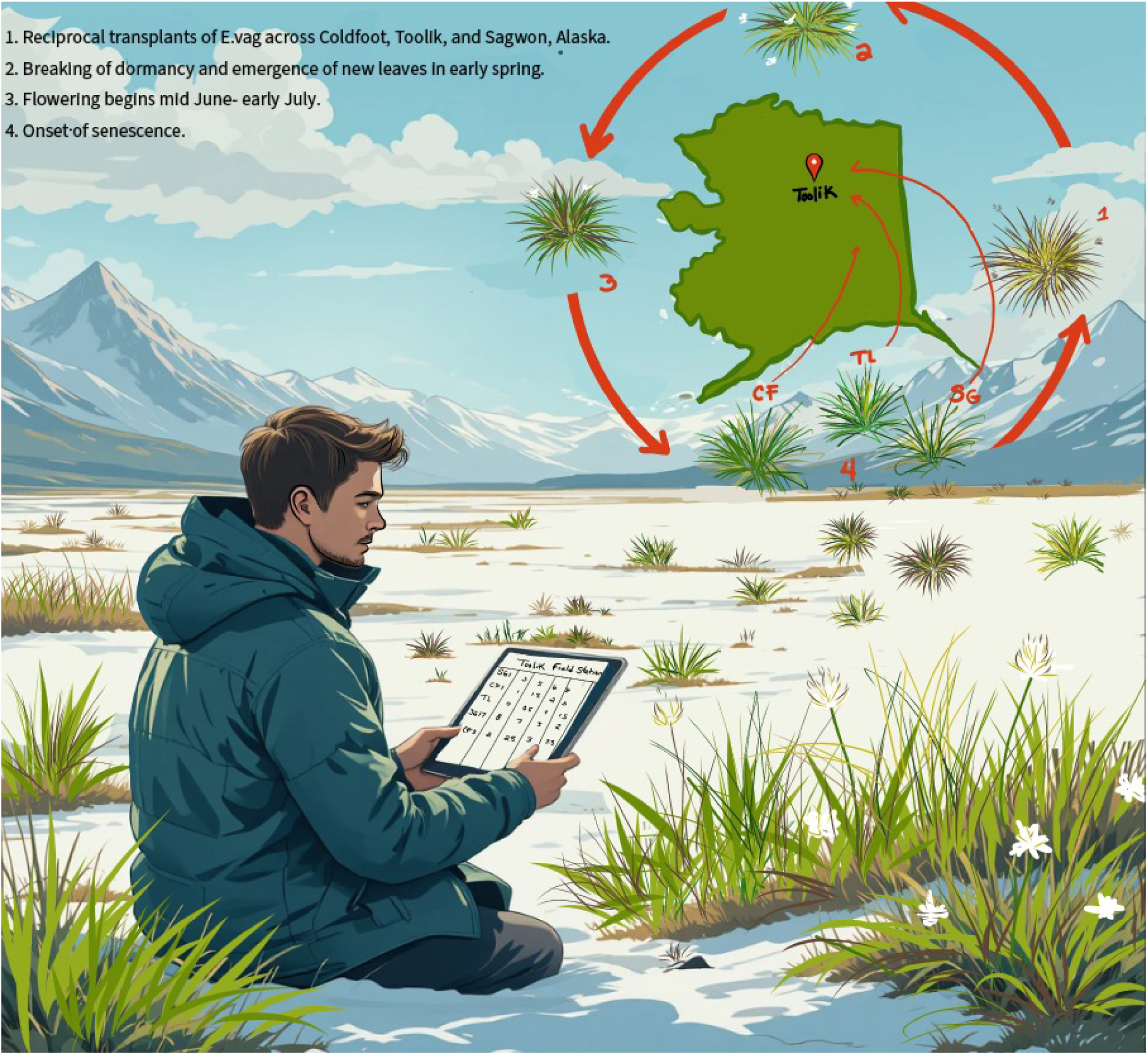

## Introduction

At Toolik Field Station (TFS), research on ecotypic variation started in 1979 when tussocks of *Eriophorum vaginatum* were brought from TFS to tussock tundra near Eagle Summit in the White Mountains of central Alaska. They survived the winter and started growing the following spring, inspiring an effort to establish a reciprocal transplant garden network in the tradition of Clausen, Keck, and Hiesey (1940), which showed that much of the variation in leaf length had a genetic basis. Five gardens were planted: Eagle Creek, No Name Creek south of the Brooks Range and Toolik Lake, Sagwon, and Prudhoe Bay north of the Brooks Range (Shaver, Fetcher, and Chapin, 1986). A sixth garden at Coldfoot was added in 1982 to increase representation of southern populations. Subsequent analyses showed that the northern ecotypes were less plastic in response to transplanting than the southern ecotypes (Fetcher and Shaver, 1990). This experiment was harvested in 2011 and showed that the different populations were adapted to their home environments demographically (Bennington *et al*., 2012) and physiologically (Souther *et al*., 2014). Subsequent analyses revealed that the optimum habitat for the populations had shifted northwards of their present locations, possibly as result of warming induced by climate change (McGraw *et al*., 2015).

In 1983 at Eagle Creek, a tussock from Toolik Lake was observed to senesce much earlier than the local tussocks. This observation led to a project to determine whether there were significant differences in phenology between ecotypes and whether these differences could influence how the populations would respond to climate change. A common garden experiment with all six populations was established at Toolik Lake in 2011 (Parker *et al*., 2017). To investigate how the different ecotypes would respond to arctic warming, open-top chambers were used to warm half of the transplants. A second reciprocal transplant garden was planted in 2014 at Coldfoot, Toolik Lake, and Sagwon and open-top chambers were again used to cover half of the transplants at Toolik Lake and Sagwon (Parker *et al*., 2021). The transplanted populations maintained patterns of phenology similar to those that they had at their home sites. Warming increased growth but produced little effect on phenology (Parker *et al*., 2017; Parker *et al*., 2021), indicating that a factor other than temperature might be driving this *E. vaginatum*’s phenology.

Global warming is causing the growing season to expand in high latitudes, exposing native plants to new environmental challenges and competitive landscapes. The increased variation in temperature associated with climate change may increase the risk of frost damage during the shoulder seasons (Augspurger, 2013; Liu *et al*., 2018). This risk might become especially severe in the Arctic, where the climate is warming three times faster than the rest of the globe (Zhou, Leung and Lu, 2024). Considering that plants in this region already have to cope with the frequent occurrence of summer freezes (McGraw, Chester and Stuart, 1983), the selective pressure for freeze-avoidance mechanisms only tends to grow in strength. Concomitantly, a lengthened growing season offers additional growth opportunities for plants that have evolved to use extremely short growing seasons. A longer growing season can also change competition dynamics among plants, with plants that manage to remain active for most of the summer having a potential selective advantage (Mulder and Spellman, 2019). These two opposing selective forces create a tradeoff between maximizing growth over an expanding season and avoiding frost damage at the shoulder seasons (Spring and Fall), which in turn defines an axis of phenological strategies. On one hand, freezes select for plants that leaf-out later and senesce earlier, thereby avoiding frost damage at the shoulder seasons, i.e. a conservative phenological strategy (CaraDonna and Bain, 2016). On the other hand, plants that remain active longer can grow more and become stronger competitors, i.e. an acquisitive strategy (Chen *et al*., 2020).

To predict the effects of climate change on arctic ecosystems, we must first understand the mechanisms that drive phenological responses in the plants of the region (Cleland *et al*., 2007). Experimental warming advances leaf-out and delays leaf senescence across multiple arctic sites (Collins *et al*., 2021), but warming effects on leaf senescence in the Arctic are not always clear and may be year-, site-, and species- specific (Oberbauer *et al*., 2013; Khorsand Rosa *et al*., 2015; Chen et al., 2020; Parker *et al*., 2021). In fact, leaf senescence of tundra plants has been substantially understudied compared to leaf-out (Prevéy *et al*., 2022). While low temperatures can be a primary driver of leaf senescence, light is also known to play a critical role (Liebsch and Keech, 2016; Woo *et al*., 2019; Jiang *et al*., 2022). Indeed, seasonal changes in day length (photoperiod) have been shown to be most critical for the timing of senescence in high latitudes (Gill *et al*., 2015). Plants can perceive changes in light across a broad spectrum through photoreceptors which regulate a broad range of physiological and phenological processes (Li, Liang and Liu, 2022). While all photoreceptors appear to be involved in some form of temperature response, some have been shown to function as thermosensors (Li, Liang and Liu, 2022). For example, phytochrome conformation along with other metabolites interact with external stimuli such as temperature to tightly control leaf senescence (Sakuraba, 2021). How the balance of light and temperature cues affect senescence in the Arctic, where summers are marked by weeks-long periods of continuous sunlight, remains unknown. Are there specific hours of darkness, temperature thresholds, or a combination of both that will trigger senescence in arctic plants?

The temperature and light thresholds that trigger senescence have likely evolved to protect plants from frost damage (Cleland *et al*., 2007; Bernhardt *et al*., 2020; Neuner *et al*., 2020). If triggered by a prolonged period of low, but above-freezing temperatures, for example, senescence may help plants avoid frost altogether, but this strategy alone is insufficient. Mid-summer freezes are not uncommon in the Arctic and could cause premature senescence if plants relied solely on temperature cues (McGraw, Chester and Stuart, 1983). It is thus likely that arctic plants have evolved to anticipate the end of the growing season by using light cues, specifically, the return of dark night hours (Shaver and Kummerow, 1992). Unlike the chaotic nature of temperature regimes, seasonal changes in day length are nearly perfectly predictable, with changes measured in milliseconds per century (Shahvandi *et al*., 2024). As long as light and temperature remain tightly coupled, plants can rely on day length to avoid frost damage (Bernhardt *et al*., 2020). However, when this coupling shifts across space or through time due to climate change (Ettinger *et al*., 2021), the more competitive plants may be those that are less rigid in photoperiod thresholds that trigger senescence or those who can track their optimum climates through migration.

Light and temperature regimes vary significantly across the Arctic, even over relatively short distances. For instance, the closer one gets to the poles, the more rapidly the day length regime changes, with faster transitions between 24-hour daylight and 24-hour darkness (Taulavuori, Sarala and Taulavuori, 2010). Temperature regimes also vary spatially due to latitude, elevation, and climate barriers, like mountain ranges. Because arctic species are so geographically widespread, they are exposed to diverse combinations of seasonal light and temperature regimes across their distribution range (Shaver, Fetcher and Chapin, 1986). Variation in local environmental conditions, in turn, may cause local populations to evolve site-specific light and/or temperature thresholds for their phenological responses (Clausen, Keck and Hiesey, 1940; Partanen, Koski and Hänninen, 1998; Peterson, Doak and Morris, 2018; Parker *et al*., 2021). Local adaptation thus leads to the formation of distinct ecotypes, which are “distinguished primarily by their reaction to the environment” (Stebbins, 1950). While ecotypic diversity allows species to thrive across wide distribution ranges, it may also increase their vulnerability to climate change (Peterson, Doak and Morris, 2018).

Local populations adapted to responding to site-specific day length cues might be at risk due to the temporal realignment between light and temperature regimes caused by climate change (Bernhardt et al., 2020; Ettinger *et al*., 2021). In the Arctic, this is most notably seen at the end of the summer, with above freezing temperatures persisting well after the 24-hour daylight period ends. These changes have accelerated in recent years: the highest temperatures on record for Northern Alaska, e.g., occurred in August 2024 (Ballinger *et al*., 2024). If ecotypes are adapted to senesce at a particular photoperiod threshold, they may not be able to take advantage of the expanding growing seasons (Shaver and Kummerow, 1992) and thus be at risk of being competitively displaced (Chen *et al*., 2020). Competitive dynamics among plants are now affected in arctic habitats due to climatic and other anthropogenic change. In interior Alaska, e.g., introduced species can senesce much later in autumn than native species, potentially giving the invaders a competitive advantage over the residents (Mulder and Spellman, 2019). The threat of competitive displacement comes also from sub-arctic species encroaching from the south (Myers-Smith and Hik, 2018) and increasing dominance of co-occurring shrubs (García Criado *et al*., 2020; Hollister *et al*., 2015). These threats are exacerbated by gene flow barriers such as mountain ranges and treelines that can prevent the migration of potentially better adapted ecotypes from south to north (Stunz *et al*., 2022, McGraw *et al*., 2015). Given the rapid pace of climate change, the growing encroachment of invading species and shrubs in arctic ecosystems, coupled with limits to northward migration, it seems increasingly unlikely that native ecotypes will have enough time to adapt phenological responses appropriate to their changing environment (Jump and Penuelas 2005). This might be the case for *Eriophorum vaginatum* L. (Cyperaceae), a foundation species of many Arctic ecosystems and an excellent model for studying adaptation to local climates.

Tussock cottongrass (*E. vaginatum*) is a widespread species, typical of moist, acidic tundra, but ranging from permafrost bogs in the taiga to alpine tundra in the top of mountains in lower latitudes to the coastal plains of the Arctic Ocean (Wein 1973). *Eriophorum vaginatum* can form distinct ecotypes across ecosystems and habitats, something that has been demonstrated by both ecological (Shaver, Fetcher and Chapin, 1986; Fetcher and Shaver 1990; Bennington *et al*., 2012; Parker *et al*., 2017; Parker *et al*., 2021; Curasi *et al*., 2019) and genetic (Stunz *et al*., 2022; Mohl *et al*., 2020) evidence. In many of the ecosystems where it occurs, *Eriophorum* is the dominant species contributing significantly to carbon storage. With temporary roots that regrow annually, these plants function like carbon pumps, drawing carbon from the atmosphere and storing it belowground (Curasi *et al*., 2022). However, with increasing permafrost depth and the expansion of shrubs in the tussock tundra, stored carbon is increasingly being released, thus converting the tundra from a sink to a source of carbon emission (Knoblauch *et al*., 2021). Accurately predicting the fate of *E. vaginatum* is therefore critical for the conservation of arctic ecosystems and their vast carbon stocks (Curasi *et al*., 2022).

To assess the adaptability of *E. vaginatum* to a rapidly warming Arctic, we used a reciprocal transplant experiment established in 2014 along a latitudinal gradient transecting tundra and taiga ecosystems in Alaska (Schedlbauer *et al*., 2018; Parker *et al*., 2021). Across this gradient, both temperature and light regimes were expected to vary by a substantial degree, so we first detailed and compared the climates at each common garden. To examine the extent to which ecotypes rely on the coupling of temperature and light cues, we monitored plant vegetative phenology for three summers (2022-2024). Using a new machine learning algorithm, we were able to estimate the timing of the onset of leaf senescence for almost all monitored plants. Lastly, we used a Bayesian framework to model the differences in phenological responses across ecotypes and common gardens, as well as the light and temperature conditions by the time plants started to senesce. This approach allowed us to better speculate on the fate of this keystone species amid current threats posed by climate change and increased competition.

### Conceptual framework

As the Arctic warms up, plants are facing new challenges with ever expanding potential growing seasons. Once well adapted to their local climates, these plants are now being even more strained by two major, but opposing, selective forces at the end of the growing season: frost-damage and competition. In this environmental scenario, plants that can remain active longer at the end of the summer (acquisitive strategy) may get the upper hand against their competing neighbors, but they do so at the cost of facing higher risk of freezing compared to plants that senesce before the winter begins (conservative strategy). To test the extent to which these selective forces have shaped the phenological responses of arctic plants, we tracked the onset of leaf senescence for the plants used in a decade-long reciprocal transplant experiment. Previous research using the same reciprocal transplant garden (Parker *et al*., 2021) found that ecotypes from the south senesced later when moved north, but northern ecotypes retained earlier senescence when moved south and, in general, *E. vaginatum* ecotypes retained similar growing season length of their homesites. Because these sites lie along a latitudinal gradient, the experiment allowed the comparison of plant responses under contrasting day length regimes. While this allowed a measure of ecotype specific response, these studies did not attempt to explicitly relate variation in senescence timing to the relative roles of changes in temperature and light in triggering plant senescence, i.e., the mechanistic basis of phenological responses. Moreover, these previous studies were conducted in the years following transplantation, i.e., while plants were likely still acclimating to their new environment. In this new study, we thus addressed the following questions: 1) How different are the climates across these three sites, especially regarding the length of the growing season and the risk of summer freezes? 2) How are climate and light regimes coupled in each of these sites? 3) Do *E. vaginatum* ecotypes still retain site-specific phenological responses after 8-10 years of being transplanted? 4) Are these ecotypes responding to temperature and/or light cues?

Independent of climate differences across sites, the simplest hypothesis would be for all plants, no matter what source population they come from, to respond to a single temperature threshold above freezing (H_0_). This ultra conservative strategy would help plants prevent frost-damage no matter where along the latitudinal gradient they find themselves. Such a strategy would not require local adaptation, but would prove to be competitively disadvantageous in sites where the probability of mid-summer freezes is not negligible, causing plants to senesce too early during the growing season. Previous results (Parker *et al*., 2021) clearly preclude this scenario, but there is still a chance that after a long period of acclimation plants would converge towards a single temperature threshold.

Alternatively, and given that the summer ends earlier at higher latitudes, one could hypothesize that northern populations have been selected to be more cold hardy than their southern counterparts. If that is the case, then northern plants’ senescence should be triggered by a lower temperature threshold compared to southern plants (H_1_), thus allowing northern plants to better exploit their shorter growing seasons. A variable temperature threshold would thus likely improve the competitive ability of this species across its latitudinal range, albeit at the cost of higher freeze exposure. Although the risk of frost damage may be minimized e.g. by the production of frost-resistant tissues (CaraDonna and Bain, 2016), temperature thresholds alone can still not protect plants from the great year-to-year fluctuation in temperature regimes that may happen within a single site. This is especially the case if higher average temperatures come with higher temperature fluctuation. For example, during a warmer than average year with mid-summer freeze events, plants responding to a given temperature threshold would senesce prematurely while their competitors enjoy a prolonged growing season. In fact, this hypothesis has also not been backed by preliminary work on this experiment, which suggested that temperature alone does not drive senescence (Parker *et al*., 2021). Again, these results might have been different, had the plants been given enough time to acclimate to their new locations, and they do not preclude that temperature may still have a supporting role in driving senescence.

A hypothesis that is more compatible with previous findings (Parker *et al*., 2021) is that this species’ phenology responds to day length thresholds instead. Indeed, photoperiod is a reliable and common trigger for senescence, especially at northernmost latitudes (Gill *et al*., 2015). Since day length in the Arctic is dependent on latitude, with the 24-h daylight period ending later farther north, it can be hypothesized that northern populations would respond to shortening day lengths (or longer nights) more strongly than those in the south (H_2_). This is a more complex phenological strategy, as it requires both local adaptation and a latitudinally varying light threshold, but it could relax the trade-off between avoiding freezes and exploiting the potential growing season. By combining acquisitive and conservative elements, this strategy may allow plants to: 1) more fully exploit their growing seasons by ignoring mid-summer freezes (acquisitive feature); and 2) escape frost damage, whose risk during the late growing season may be concealed by above-average warm temperatures (conservative feature).

Of course, plants could have an even more complex strategy, with light and temperature thresholds that were both site-specific (H_3_). For example, plants’ temperature thresholds could be dependent on the photoperiod threshold, thus time-calibrating their assessment of the risk of freezes. Shifting from a risk-prone (acquisitive) to a risk-averse (conservative) strategy along the growing season would enable plants to safely exploit good and bad years alike: they could still dismiss the risk of mid-summer freezes while remaining active for longer during an exceptionally warm fall. In fact, both day length and temperature seem to be driving senescence throughout the northern hemisphere (Gill *et al*., 2015). It is not clear though whether the combined light-temperature thresholds of senescence vary across local populations, as it has been documented for bud burst in Norway spruce (Partanen, Koski and Hänninen, 1998).

Within the context of our reciprocal transplant experiment, we can make the following predictions regarding the onset of leaf senescence:

H_0_: If temperature is the primary driver of senescence and plants respond to the same temperature threshold, then: 1) all plants transplanted into a common garden should senesce at the same time, regardless of their site of origin (Fig.1); and 2) plants transplanted north should senesce earlier than they do in their home sites.

**Figure 1.**
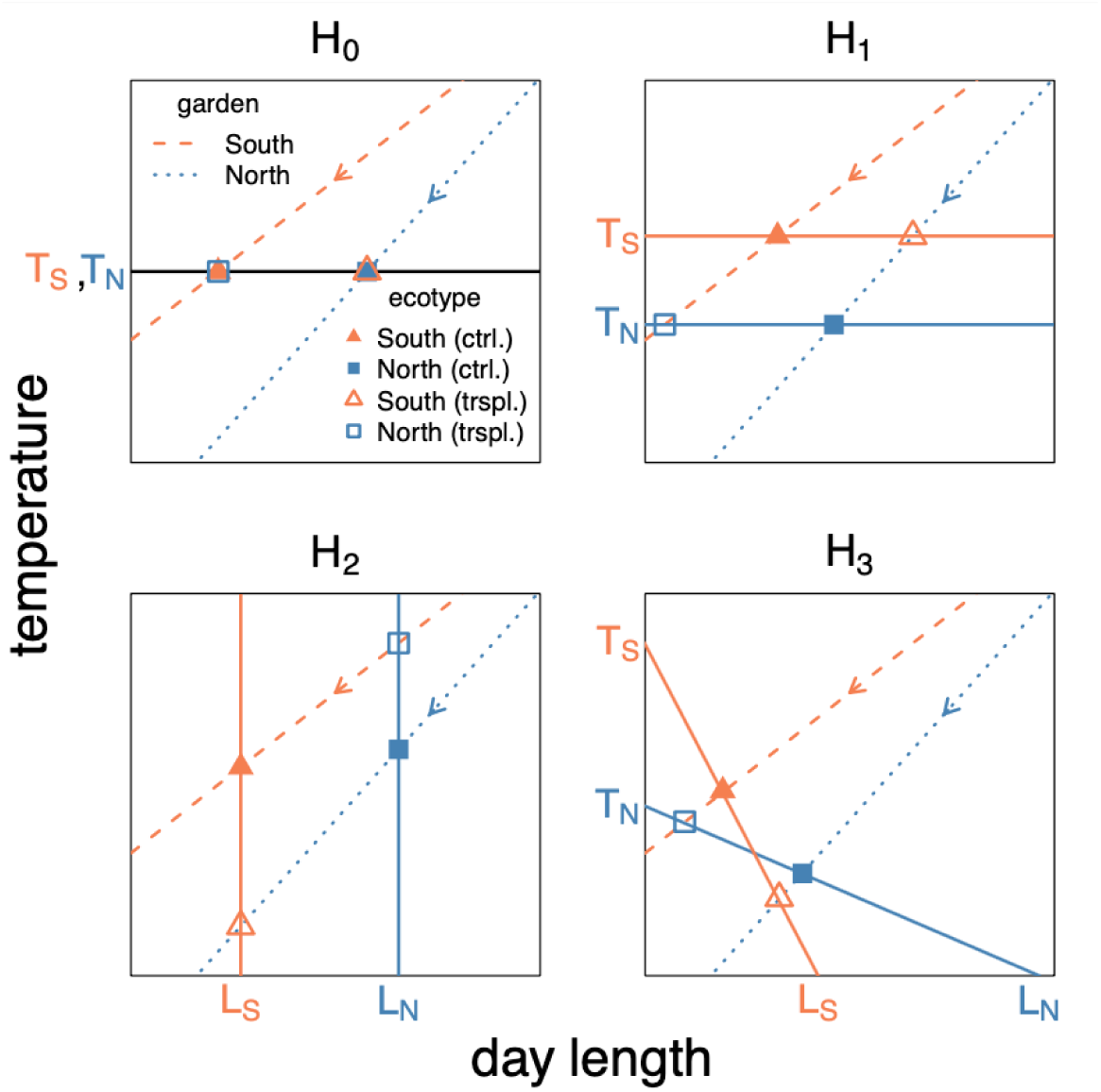
Graphical conceptualization of the four hypotheses (H_0_-H_3_) outlined by the conceptual framework of this study. Habitat curves (traced and dotted lines) represent the natural reduction in temperature and day length through the growing season after the summer solstice in hypothetical sites in the south and in the north, respectively. The arrowheads in these lines show the direction time flows: from long and warm days (upper right corner) to cold, short days (lower left corner). Ecotypic response curves (continuous lines) represent hypothetical temperature and day length thresholds that trigger leaf senescence for southern (S) and northern (N) ecotypes. The points where habitat and response curves cross (squares and triangles) mark the moment when plants’ phenological thresholds have been met, thus representing the onset of leaf senescence for ecotypes in their native habitats (filled points) and when transplanted across sites (empty points).

H_1_: If temperature thresholds decrease with latitude, then: 1) northern ecotypes should senesce later than their southern counterparts across all common gardens (Fig.1); and 2) all plants moved south should senesce later than they do in their site of origin.

H_2_: If plants respond instead to day length thresholds, and these thresholds increase with latitude, then: 1) southern ecotypes should senesce later than their northern counterparts in each and all common gardens (Fig.1); and 2) all plants moved north should senesce later than they do in their site of origin.

H_3_: If both H_1_ and H_2_ are true, but temperature thresholds also increase as days become shorter, then the temporal order by which ecotypes start to senesce could switch across the gardens (Fig.1), i.e., there should be an interaction effect between ecotype and garden. Making predictions more precise than this would require us to know how excatly ecotypes’ temperature threshold changes with shrinking day lengths. Nevertheless, a possible outcome, unique to H_3_, would be for southern plants to senesce later than northern ones in the north, while northern plants senesce later than southern ones in the south.

## Materials and methods

### Study sites and experimental design

To test how well adapted *E. vaginatum* populations are to their local climate, we used a long term reciprocal transplant experiment in Northern Alaska. The three common gardens used were established in 2014 (Parker *et al*., 2021), one to the south and two to the north of the Brooks Range (Fig.2). The southernmost site is located next to Coldfoot Camp (CF; 67.26°N, 150.17°W, 335 m.a.s.l.) in muskeg habitat encroached by black spruce, *Picea mariana (Mill.) Britton, Sterns & Poggenb.* The northernmost garden is in a site of typical tussock tundra (SG; 69.42°N, 148.71°W, 293 m.a.s.l.). In between, close to Toolik Lake Field Station (TL; 68.63°N, 149.58°W, 762 m.a.s.l.), lies the second tussock tundra garden mixed with dry heath patches. Although the latitudinal contrast between CF and SG (2.16°) is only a small fraction of this species’ range, the sites are found in two different ecosystems, tundra to the north and taiga to the south, and climate differs drastically between the gardens split by the Brooks Range, with CF being much warmer than TL and SG (see results). In each garden, plants brought from the other two sites, as well as local plants, were transplanted into plots with three plants each, being all three plants within a plot of the same origin (Fig.2). These transplants were done at the same point where a local plant existed before to avoid placing plants in sites unsuitable for *E. vaginatum*; the local plants used in the experiment were simply moved around to serve as controls. In total, there were 216 plants distributed into three common gardens, with 8 plots per ecotype, containing three plants each (Fig.2). The TL and SG gardens also included a warming treatment – data from these plants are not included here as they are outside the scope of this paper. We also excluded plants from 12 plots at TL that were subjected to a 2022 daylength manipulation pilot experiment that year.

**Figure 2.**
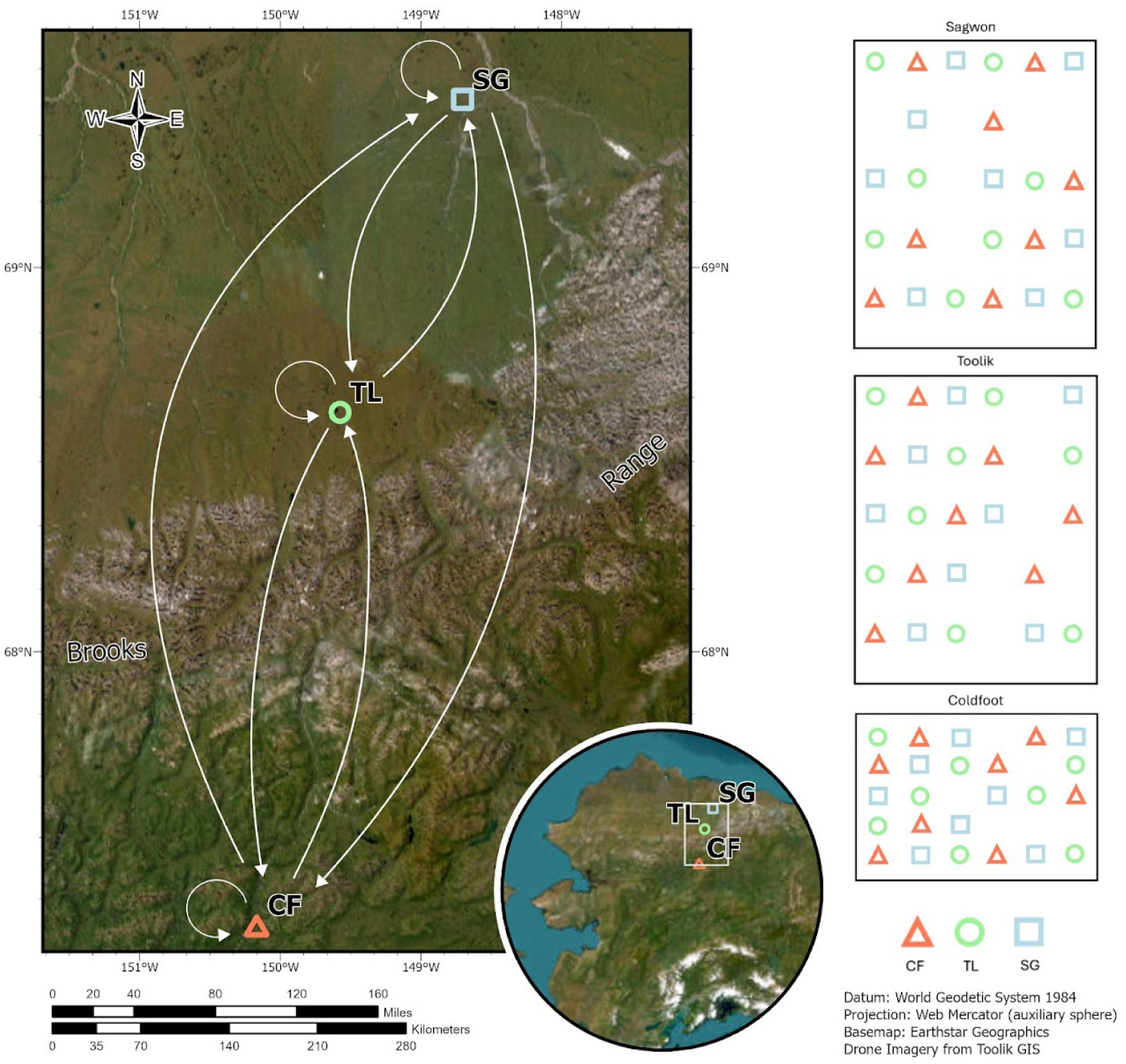
Map showing the location in Northern Alaska (inset) and across the Brooks Range (left) of the three common gardens used in this study: Coldfoot (CF; taiga), and Toolik (TL; tundra) and Sagwon (SG; tundra). The diagrams on the right display the spatial arrangement of plots in each reciprocal transplant garden, each plot having three individuals of *Eriophorum vaginatum*.

### Defining the potential growing season

To define the start and end of each year’s potential growing season (PGS) we used snow depth and air temperature data (Fig.3) from weather stations located close to each of the three gardens. Although *E. vaginatum* can start growing under the snow, this only happens a few days before snowmelt (Starr and Oberbauer, 2003), so we considered the start of the season (SOS) to be when snow had completely melted and the end of the season (EOS) to be when snow permanently (for that year) covered the ground once again. We then calculated the length of the season (LOS) in each site and year as the difference between SOS and EOS. However, because these sites are subjected to mid-summer frosts, we also calculated the length of the PGS as the number of hours between SOS and EOS during which air temperatures remained above 0°C. The hourly air temperature and snow depth data we used were collected by the Snow Telemetry (SNOTEL) Network in Coldfoot and Sagwon (site numbers 958 and 1183, respectively; United States Department of Agriculture, 2024). In Toolik, we used similar data provided by the Toolik Field Station (TFS) (Environmental Data Center Team, 2025), combined with data derived from phenocams. Snow depth has only been monitored at TFS since 2019, so we used images from a time-lapse camera mounted in the moist acidic tundra since 2012 (https://www.uaf.edu/toolik/edc/monitoring/abiotic/time-lapse.php) to estimate the time of complete snow melt and cover. Because the weather stations and phenocams are not installed right next to our gardens, and snow cover patterns may vary spatially, we validated our estimates by visually inspecting the images taken by the two Sentinel-2 satellites (https://browser.dataspace.copernicus.eu). Although this database has only a 5-day temporal resolution, the images could still confirm our snow cover estimates. Finally, because each weather station started operating at a different time, we used the period (2012-2024) for which we had the same information for all three sites – except for 2014 in Sagwon, the record is complete for all sites and years within this period. With PGS defined for each site and year, we could then characterize summer freeze and light regimes for each of our gardens.

**Figure 3.**
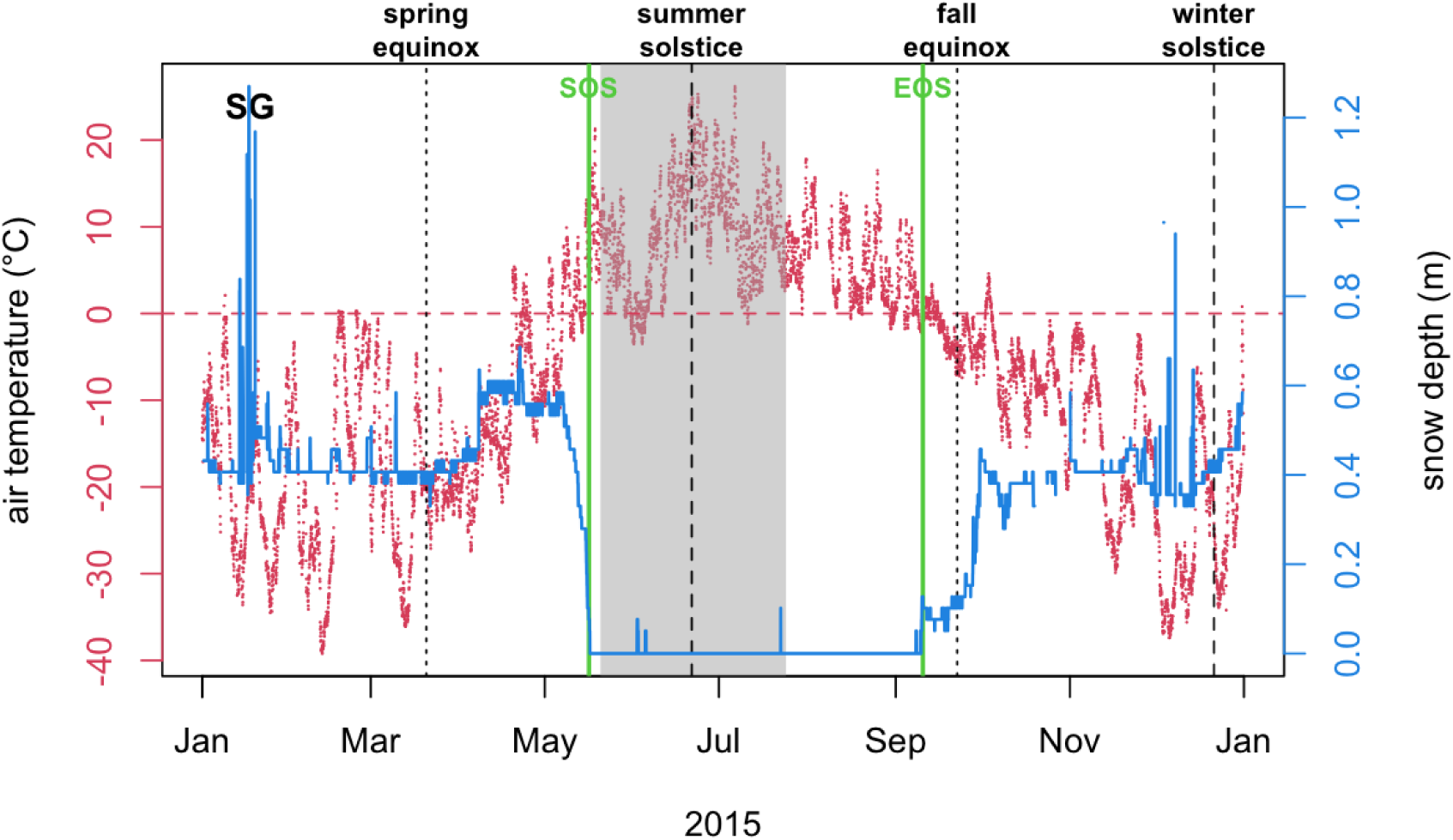
Hourly air temperature (red points) and snow depth (blue, continuous line) in 2015 at Sagwon, shown here as an example of how the potential growing season (PGS) was calculated. Snowmelt in the spring and snowcover in the fall mark the start and end of the PGS (SOS and EOS, respectively; vertical green lines). Also shown are the main light regime landmarks: equinoxes (12-hour day length) and the 24-hour day length period (grey area) around the summer solstice.

### Quantifying summer freeze and light regimes

To compare summer climate across all three sites, we calculated a series of metrics on the freezing events that took place in each site and year during the pre-defined PGS. The simplest measure is freeze frequency, i.e., the number of freeze events that took place during each PGS (freeze events). Each of these events was defined as an uninterrupted sequence of hourly data during which air temperatures (T_air_) were at or below 0°C. Next, we calculated the intensity of each freeze event as both the mean and minimum T_air_ reached and the cumulative negative degree hours during each event. We also calculated the duration (in hours) and timing (mean day of the year) of each event. Lastly, to describe the potential interplay between freeze and light regimes, we crossed T_air_ with hourly solar elevation angle (SEA) data derived from the series of equations provided by the National Oceanic and Atmospheric Administration (https://gml.noaa.gov/grad/solcalc/calcdetails.html). We established two metrics: 1) the proportion of freeze hours that took place while the sun was at or below the horizon (SEA < 0°); and 2) the cumulative positive degree hours ahead of each freeze hour until the end of the PGS.

### Monitoring leaf phenology

To track vegetative phenology of *Eriophorum*, we generally followed the method of choice for this species (Shaver and Laundre, 1997; Parker *et al*., 2017). An important difference is that we measured only the green portion of each leaf, ignoring the brown, dried tips (typical of the oldest leaves formed in the previous summer) and the yellow portions of senescing or dying leaves during the fall. Roughly every week, we measured all transplanted plants in each of the three common gardens during the summers of 2022-2024. We started monitoring the plants’ leaf green length after snow melt and continued monitoring until late in the growing season (Tab.S1). The monitoring frequency within the season on average varied between 7 and 9 days (Tab.S1). During each visit, the green length of all the leaves in 1-3 tillers per tussock was measured for all the 216 transplanted plants, totalling 555 tillers, 3,182 leaves, and 33,060 measurements. Then for each monitored date, we summed together the length of all leaves in each tiller and used it to detect the onset of senescence as described in the next section.

### Detecting the onset of leaf senescence manually

We structured our analysis to determine the crucial time when plants are triggered to start senescing in preparation for the winter. To find this elusive turning point, we first plotted our weekly data on total leaf green length, yielding green-up and green-down curves for each monitored tiller (Fig. 4). We then manually picked the day of the year (DOY) when tillers went from greening-up to greening-down, i.e., the onset of leaf senescence (OLS). There was not always a single peak in greenness; oftentimes the leaf phenology curve declined then increased again, or gradually declined before plateauing and then reaching a steep decline (Fig.4). To deal with this complexity, we first located the final peak greenness in the curve and selected the last DOY before greenness reached a steep decline (Fig.4). We discarded the observations for the few instances when tillers did not show an obvious peak in greenness, i.e., the curve was either monotonic, U-shaped, or was quite irregular, with many peaks and valleys, as these cases were often linked to sick, damaged, or dying tillers. This caused the number of plants sampled to drop from 216 to 213. When there was no obvious decline in greenness, i.e., greenness remained relatively stable (plateau) until the end of the monitoring period, we picked the DOY on which those tillers were last measured. To avoid biases, tiller ids were kept hidden and randomly shuffled prior to the process of handpicking the OLS.

**Figure 4.**
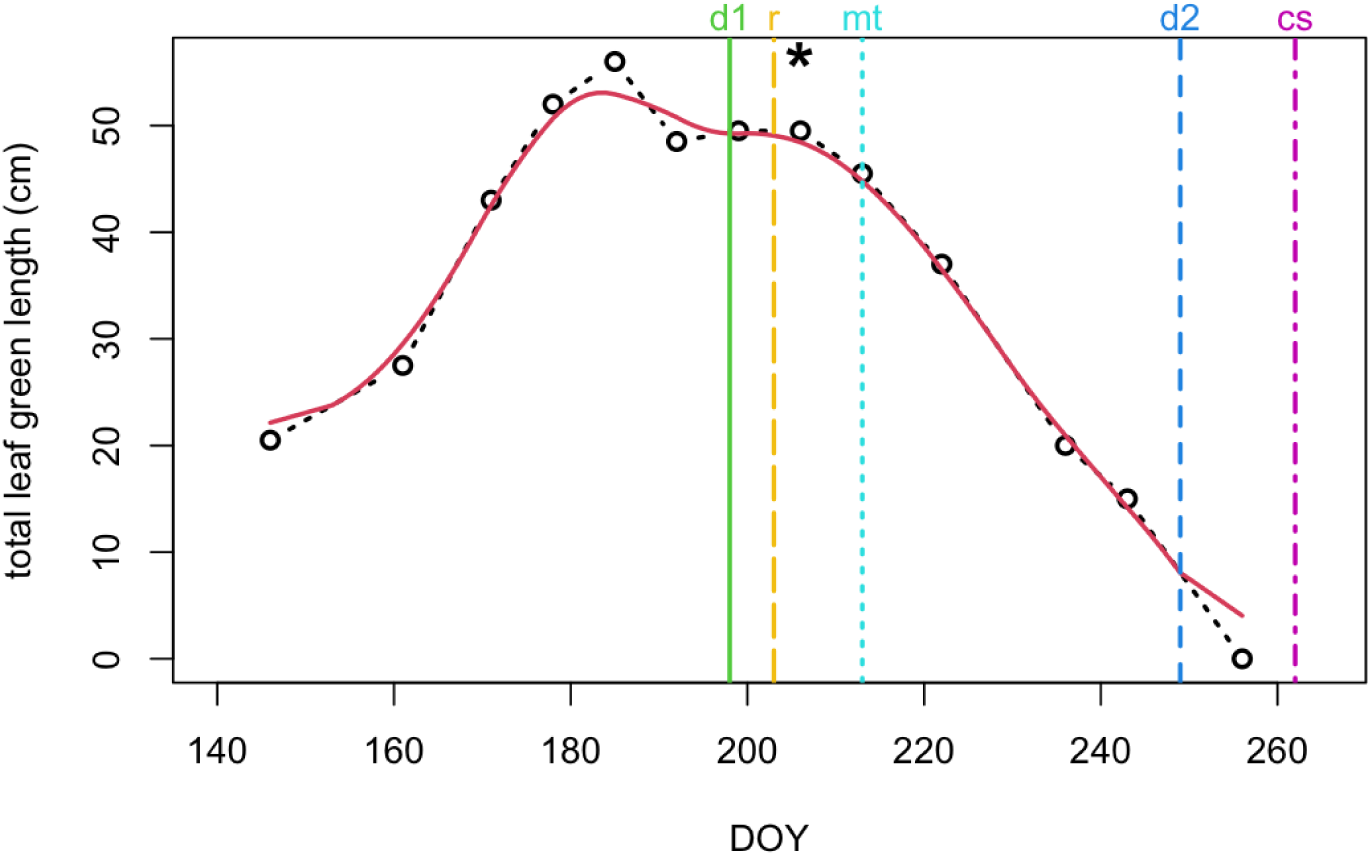
Example of an observed leaf phenology curve and the estimates for the onset of leaf senescence (OLS) for *E. vaginatum*. This example originates from a control plant in the Coldfoot Garden and shows the weekly measurements (points) of the sum of the green length portion of all leaves in a marked tiller. The black dashed-line represents the phenology curve derived from the raw data. The red continuous curve represents the smoothed total green length (rolling 14-day average) and the vertical lines represent the day of the year (DOY) of each OLS detection method’s prediction detailed in the methods (d1 = first derivative; d2 = second derivative; mt = maximum threshold; cs = cumulative sum; and r = linear regression). For comparison, we show the handpicked DOY at the OLS (asterisk).

### Estimating the onset of senescence through machine learning

To avoid other potential biases in picking the onset of senescence (OLS), we used five machine learning algorithms and cross-validated the results with our handpicked OLS (Fig.4). Starting with weekly observational data:

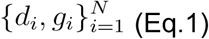

where *d_i_* and *g_i_* are the day of the year and total leaf green length, respectively, and *N* is the number of weekly measurements for each tiller. We then applied linear interpolation to obtain a daily time step:

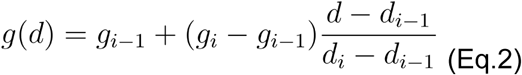

Where *i* is the index of the observation, for a given day *d* and total leaf green length *g* for that observation. Then we used a rolling 14-day average to smooth the data:

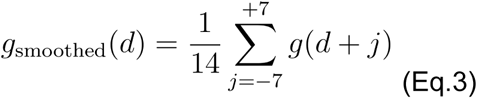

To minimize noise, leaves were excluded if they lacked measurements beyond the midpoint of the growing season, had fewer than three observations across the season, or showed a large increase in green length followed by a sharp decrease (or the opposite, a sudden decrease followed by a sharp increase), which likely indicated an error in leaf identification. The first algorithm, called *maximum drop threshold*, was computed as follows. Starting with the smoothed daily green length, we found local maxima by simply finding green lengths that are higher than their neighbors within a rolling window *W_t_*:

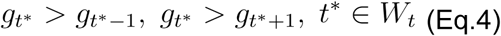

If no local peaks are found, we used the global maximum. Given a peak value *g_t_*\*, we defined a drop threshold *θ* as a fixed fraction of the peak:

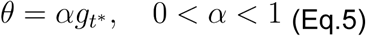

Where *α* is the threshold fraction. Using this threshold, we identified a candidate set of days after the peak where the green length falls below the threshold:

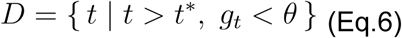

To narrow down the number of candidate days, green length needs to stay below the threshold for at least *L* consecutive days:

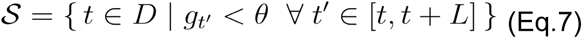

The start of senescence is the first day in this set:

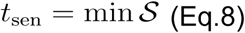

Finally we applied some constant offset Δ to calibrate the final estimated OLS to the handpicked values:

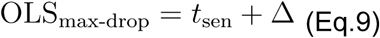

Detailed descriptions of the four other algorithms – first and second derivative, linear regression, and cumulative sum of deviation – are included in the appendix.

All parameters used in the *maximum drop threshold* (see Tab.1 for a detailed description) were optimized using gradient based descent. The loss function was constructed as the root mean squared error (RMSE) of the computed OLS compared to the handpicked OLS. We then used the OLS values produced by the algorithm with the smallest RMSE to conduct the final data analysis. Final analyses were performed in `Python` v3.12.3 with the following libraries: `NumPy` v1.26.4 (Harris *et al*., 2020) for numerical computation, pandas v2.2.2 for data handling, `SciPy` v1.14.1 (Virtanen *et al*., 2020) for optimization and statistics, and `matplotlib` v3.9.1 (Hunter, 2007) for visualization.

The four other algorithms were also tested. The *first* and *second derivative* of green length with regard to time were computed and the OLS was taken to be the first day where they were at some threshold value. Similarly, a rolling *linear regression* was applied over a certain number of days, and the day at which the slope started to become negative was used as the OLS. Finally, the *cumulative sum* of deviation from the expanding mean of green length over the season was computed and OLS was determined by when the sum increased by some value. For details on the parameters used in these other detection algorithms, see table S2, and the appendix section *Machine Learning Algorithms Detecting Onset of Leaf Senescence*.

**Table 1.**
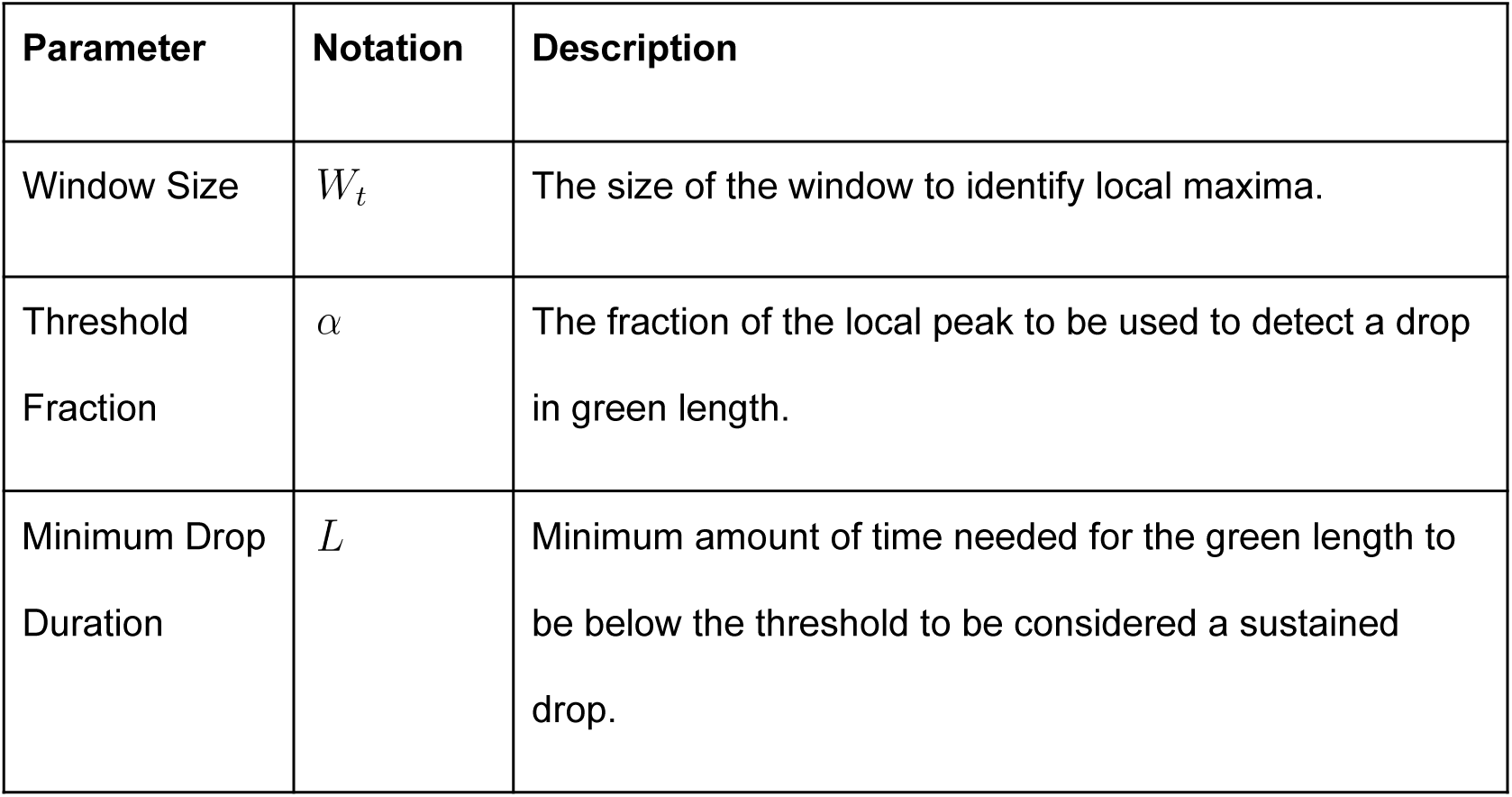
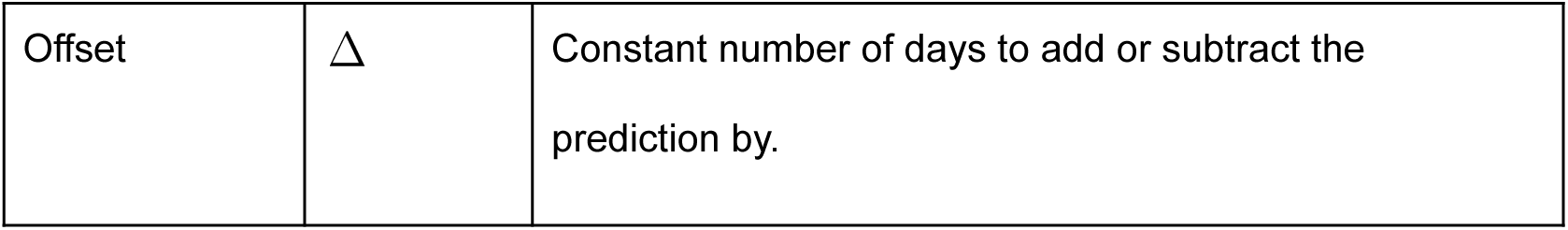
Description of the best performing algorithm (*maximum drop threshold*) used to estimate the onset of leaf senescence from leaf green length data of the monitored individuals of *Eriophorum vaginatum*.

| Parameter | Notation | Description |
|---|---|---|
| Window Size | *W<sub>t</sub>* | The size of the window to identify local maxima. |
| Threshold Fraction | *α* | The fraction of the local peak to be used to detect a drop in green length. |
| Minimum Drop Duration | *L* | Minimum amount of time needed for the green length to be below the threshold to be considered a sustained drop. |
| Offset | Δ | Constant number of days to add or subtract the prediction by. |

### Modeling the onset of senescence: effects model

To estimate how sensitive plants are to late summer freezes and seasonal changes in light, we modeled the onset of leaf senescence (OLS) – as generated by the *maximum drop threshold* model output – as a function of a plant’s population of origin (ecotype) and the site where it was transplanted (garden). To take advantage of all the information available in the data – from yearly variation in OLS for each tiller sampled all the way to overall differences across ecotypes – we used a hierarchical Bayesian framework. We first used a simple model based on the factorial design of the reciprocal transplant experiment with the aim of estimating the independent and joint effects of ecotypes and gardens on OLS. This “effects” model can be described as follows:

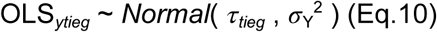

Where the identifier *y* is the year of OLS measurement, *t* is the tiller that was sampled that year, *i* is the individual plant (tussock) where that tiller belongs, *e* is the site of origin (ecotype) of that plant, and *g* is the site (common garden) where that plant was transplanted in 2014. *σ* ^2^ is thus the cross-year (2022-2024) variance in the tillers’ OLS and *τ* is the mean OLS for each sampled tiller, which was subsequently modeled as:

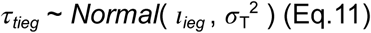

where σ^2^_T_ is the cross-tiller variance and *ι*’s are the mean OLS for each individual plant, which was then modeled as:

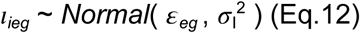

where σ^2^_I_ is the cross-plant variance and *ε*’s are the mean OLS for each ecotype-garden combination (9 total), which was then modeled as follows:

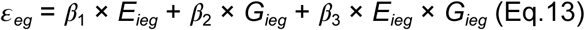

where *β*_1_, *β*_2_, and *β*_3_ are, respectively, the ecotype, the garden and the interaction effect on *ε*, and *E* and *G* are the latitudes (centered and scaled) of the plants’ sites of origin and destination, respectively. To simplify the analysis, we forced the intercept in Eq.13 to zero by centering and scaling OLS data. We could then define the prior distributions for *β* as normal distributions centered at zero and with variance of 1, while the priors for *σ* were uniform distributions constrained between 0 and 1. Parameter estimations were done using the Markov Chain Monte Carlo (MCMC) algorithm in `JAGS` through `R` using the `rjags` package (Plummer, 2023). To guarantee reliable estimates, we ran three chains simultaneously for a million iterations each, discarding the first half as burn-in and thinning the remaining to 1,000 values per chain. MCMC convergence of the main parameters were assessed through visual diagnosis of raw values, density, autocorrelation, and shrink factor plots.

### Modeling the onset of senescence: means model

For more precise estimations of the variances at each hierarchical level and the means of the nine ecotype-garden combinations, and thus complement the “effects” model, we also ran a “means” model. To do so, we replaced Eq.13 by the following:

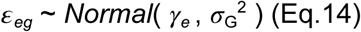

where σ^2^_G_ is the cross-garden variance and *γ* is the mean OLS for each ecotype, which was finally modeled as:

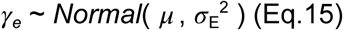

where σ^2^_E_ is the cross-garden variance and *μ* is the mean OLS for each ecotype. The prior distribution for *μ* was set as normal distribution with mean zero and variance of 1, since OLS data had been previously centered and scaled. The prior distribution for the variance terms was a Dirichlet distribution with equal weights for each of the five hierarchical levels (year, tiller, individual, garden-ecotype, and ecotype). Because the sum of all the variance parameters should add to 1, i.e., the total variation in the data, using this prior forces variance estimates to follow this condition. The simulations to estimate this model’s parameters, as well as the diagnostic analysis of MCMC convergence, were run in the same way as the effects model.

### Modeling weather conditions at the onset of senescence

Finally, to gain insight into the factors driving the onset of leaf senescence, we replaced OLS with two variables related to freeze and light regimes. The first variable was the minimum temperature (T_min_) registered for the 11-day period around each observed OLS. This temporal slack was used to accommodate our weekly monitoring schedule of leaf senescence. Likewise, for each observed OLS we also calculated the minimum solar elevation angle (SEA_min_) during this same period to gauge how much darkness the plants were experiencing by the time they started to senesce. The two models – “effects” and “means” – were run again in the same way as before, only this time using these two response variables, one at a time.

## Results

### Potential growing season and summer freeze regimes

Except for longer potential growing seasons (PGS) in the taiga site (Coldfoot), the summer freeze regime was surprisingly similar across all sites, with the most obvious differences being the onset and end of the PGS. As expected, PGS started earlier and ended later in Coldfoot, yielding a much longer PGS compared to the northern sites, especially Toolik. The extremes ranged from a PGS at Toolik lasting only 3 months to a PGS at Coldfoot lasting for almost 5 months (Fig.5). In addition to a later start, the daily freeze probability started higher in the northern sites compared to the southern site: there was a 44% chance of negative temperatures in Coldfoot by the start of the season (SOS), compared to 66% and 68% in Toolik and Sagwon, respectively (Fig.6a-c). The situation reversed by the end of the season (EOS), with Toolik and Sagwon having respectively a 74% and 76% chance of negative temperatures before snow cover, while in Coldfoot this probability was 85% (Fig.6a-c). During the PGS, air temperature was at or below 0°C only 10% of the time in Coldfoot, whereas for Toolik and Sagwon this figure was around 15%. These patterns are summarized by the remaining cumulative positive degree hours (CPDH) that are left in the year. While in Coldfoot, CPDH approached zero way before EOS, in the north this happens almost concomitantly with snow cover (Fig.6d-f). In spite of these differences, the freeze frequency, intensity, duration, and timing showed no substantial differences across sites (Fig.7).

**Figure 5.**
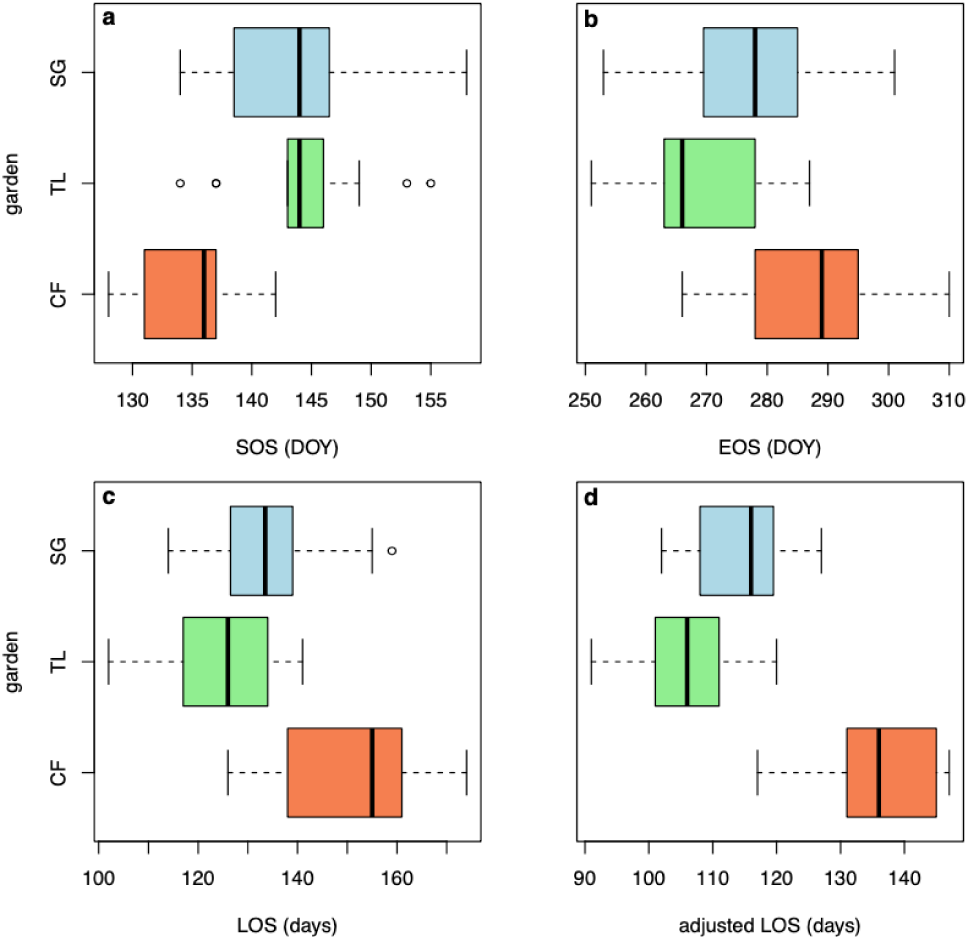
Days of the year (DOY) when the snow-free season starts (SOS; a) and ends (EOS; b) and the resulting length of the snow-free season (LOS; c) for the three studied sites: Coldfoot (CF), Toolik (TL), and Sagwon (SG). Also shown is the adjusted LOS (d), i.e., the amount of time during the snow-free season when air temperature was above freezing point. These values are based on climatic data collected between 2012 and 2024.

**Figure 6.**
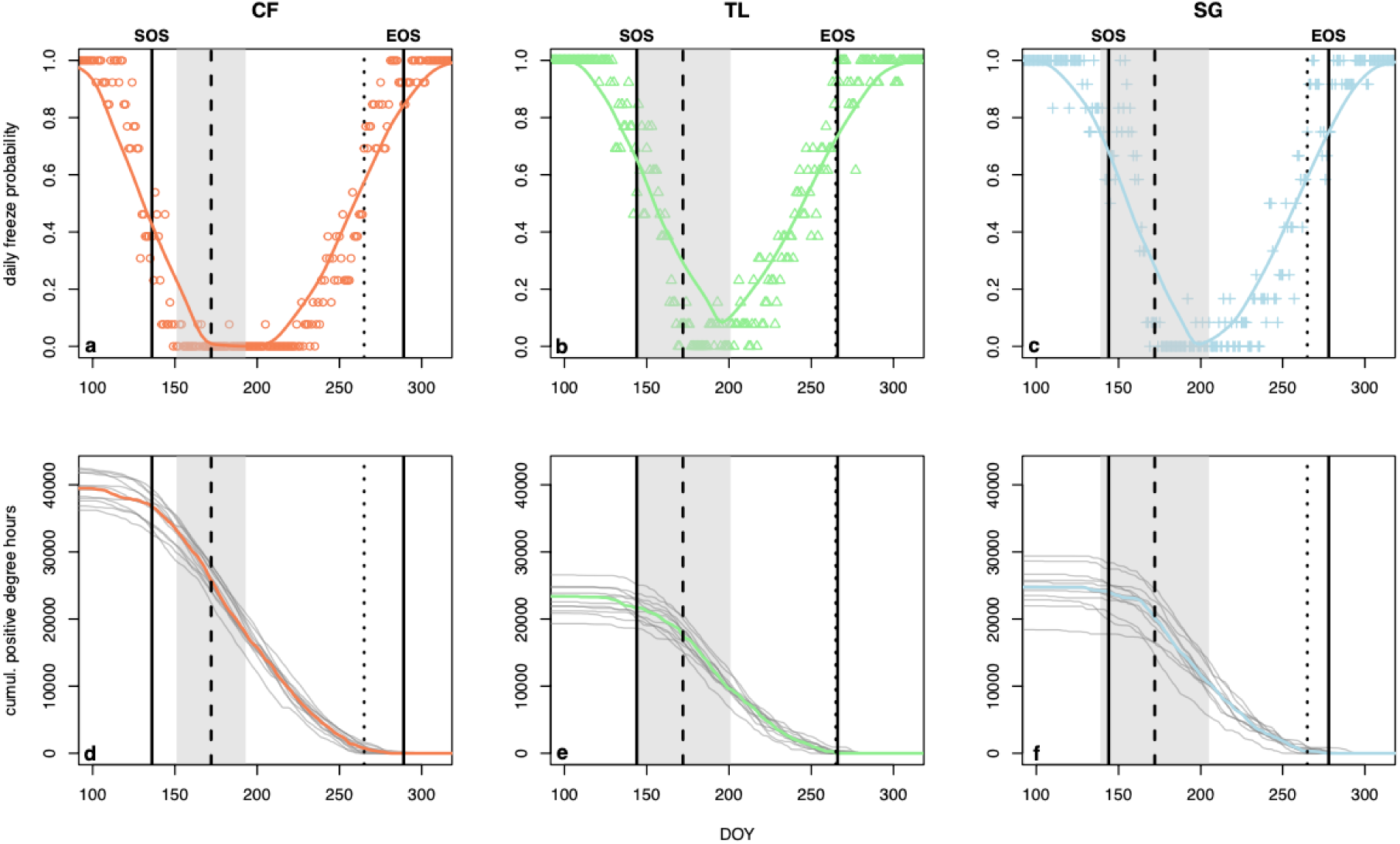
Daily probability of freezes (a-c) and cumulative positive degree hours (CPDH) ahead of each day (d-f) along the summer in each of the three study sites: Coldfoot (CF), Toolik (TL), and Sagwon (SG). These probabilities and CPDH were calculated based on hourly air temperatures collected from 2012 to 2024. Vertical solid blacklines represent the onset and end of the snow-free period (SOS and EOS, respectively). Dots in the top panels are the probability of the temperature going below freezing point for each day of the year (DOY). Grey lines in the bottom panels are the CPDH between each DOY and the EOS for each year. The continuous thick lines are locally-weighted polynomial regression of probability (top) and CPDH (bottom) by DOY. The vertical black traced line is the summer solstice and the grey area around it is the 24-hour daylight period.

**Figure 7.**
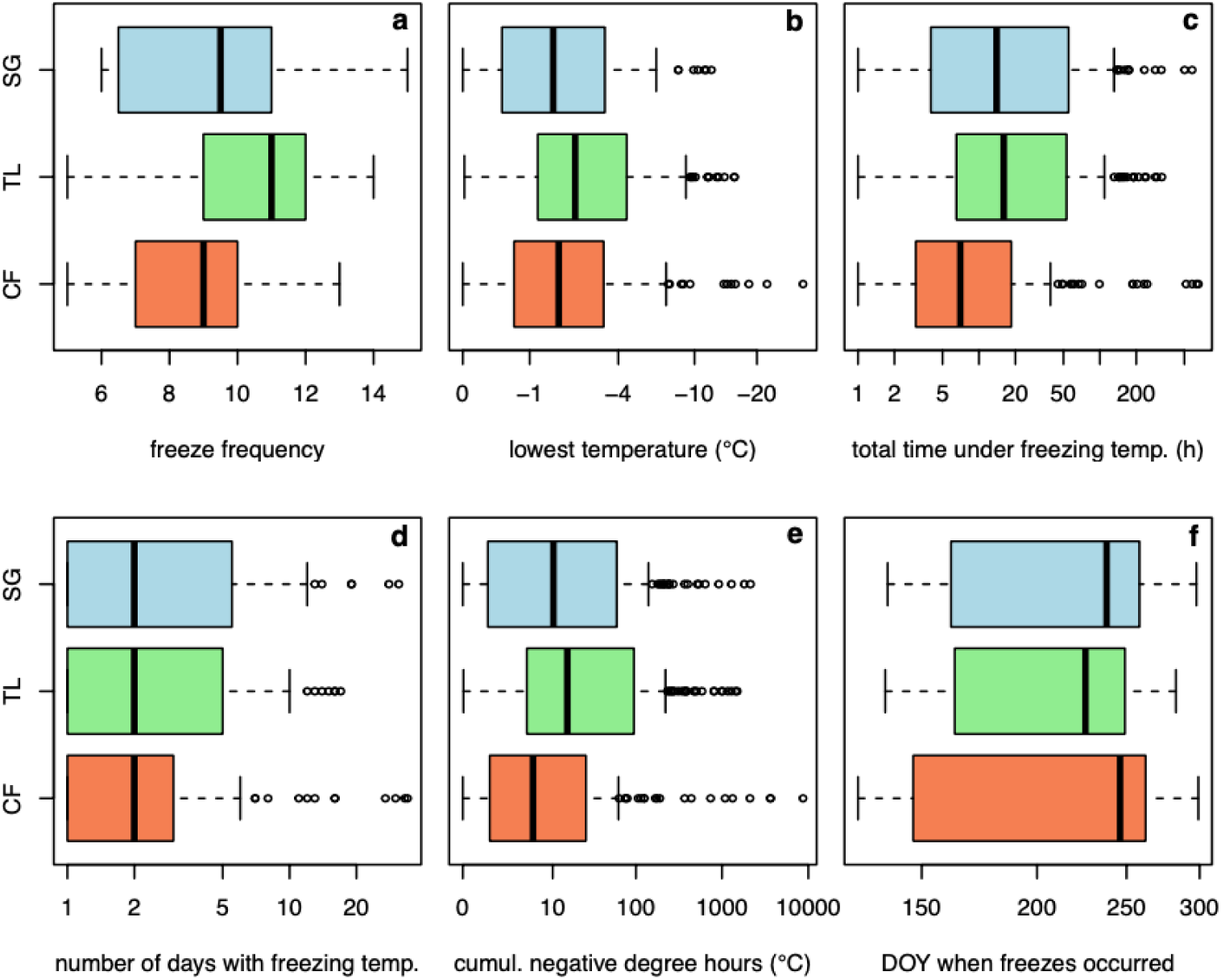
Characterization of freeze regimes during the potential growing season (PGS) across the three study sites for the 2012-2024 period. The frequency of freezing events was calculated as the number of distinct freeze events during each PGS (a). The remaining metrics were calculated for each freeze event that occurred during each PGS (b-f). These metrics reflect freeze intensity (b), duration (c-d), duration and intensity (e), and timing (f).

Each site showed a unique timing for its temperature regime. During the first part of the PGS the northern sites were equally cold compared to Coldfoot. However, Coldfoot and Sagwon tend to converge towards milder temperatures compared to Toolik towards the end of the PGS (Fig.8a,c). This might have occurred because, although Toolik has an intermediate latitude, it has the highest elevation. Lastly, both the daily freeze probability and CPDH varied predictably and monotonically with daily minimum temperatures (T_min_) after the summer solstice (Fig.8f,h). For CPDH these relationships also varied substantially across sites: by the time minimum daily temperatures dropped to 5°C, e.g., there was only a slightly higher freeze probability in Coldfoot (11%) compared to Toolik and Sagwon (9 and 6%, respectively), but a large difference in what is left of the actual growing season (15, 26, and 41% for Coldfoot, Toolik, and Sagwon, respectively). Overall, these results suggest the main climatic differences across the study sites are mostly due to how far apart the shoulder seasons are, which is when freezes are most common.

**Figure 8.**
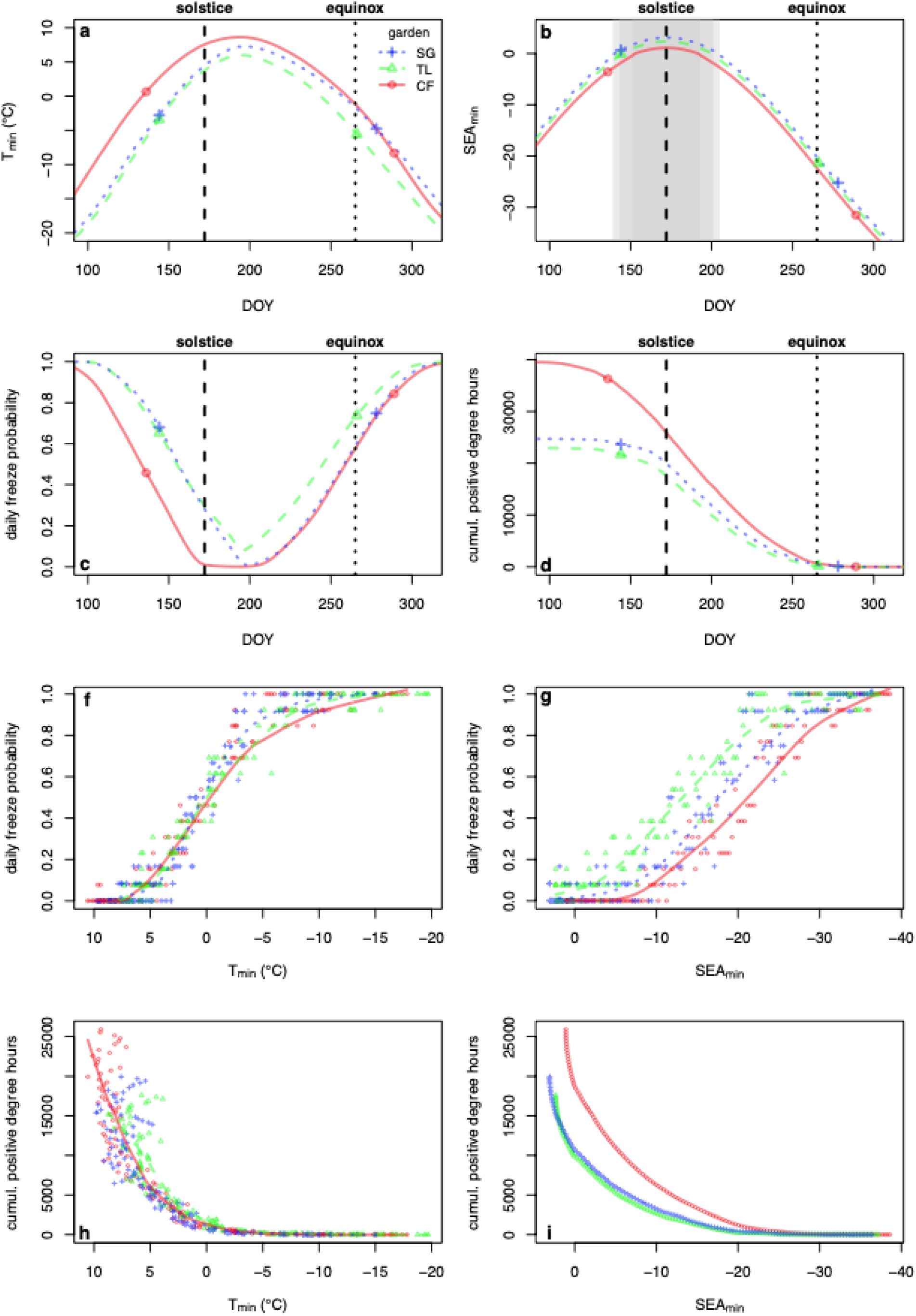
Temperature and light regimes throughout the potential growing season and how they co-vary across the study sites. Panels a-c show, respectively, temperature, light, and freeze regimes throughout the potential growing season (PGS) in Coldfoot (CF), Toolik (TL), and Sagwon (SG). Temperature regimes (a) are represented by daily minimum temperatures (T_min_) and freeze regimes (c) are represented as the probability that temperatures drop below 0°C for each day of the year (DOY). Light regimes across the study sites (b) are represented by the minimum solar elevation angle (SEA_min_) for each DOY. Panel d shows how much of the actual growing season is left along the PGS, which is represented by the cumulative positive degree hours (CPDH) between each DOY and the end of the PGS. Panels f-i show how freeze probability and CPDH after the summer solstice can be predicted from changes in T_min_ and SEA_min_. Grey areas around the summer solstice (b) represent the length of the 24-hour daylight period, which is smallest for CF and largest for SG. Except for panels b and i, all curves were smoothed by locally-weighted polynomial regressions. The start and end of the potential growing season in each site are shown as large points along these curves (a-d).

### The timing of freezes in relation to light regimes

Although the freeze regime during the potential growing season (PGS) did not vary substantially across the study sites, the timing of freeze events in relation to the light regime was unique to each site. This occurred in spite of the relatively small differences in the seasonal changes in day length across sites. The two most noticeable differences were: 1) the 24-h daylight period (24DP) lasted much longer in the northern sites compared to the southern site (42, 58, and 66 days in Coldfoot, Toolik, and Sagwon, respectively; Fig.8b); and 2) the onset of the PGS in the northern sites coincided with the start of the 24DP, while in Coldfoot the PGS started ca. two weeks prior to the onset of the 24DP (Fig.6). Although the PGS ended at different times after the spring equinox (12-hour long day) in each site, there were few cumulative positive degree hours (CPDH) left by fall equinox, with little variation across sites (Fig.6d-f and Fig.8d). In contrast, the timing of freeze events in relation to light regimes varied much more across the sites. One of the main differences is that freezes were common during the 24DP in the northern sites, but were almost absent in Coldfoot where freezes only occurred when day length had already dropped to ca. 17 h (Fig.6a-c). Another remarkable result is the distinct way the freeze probability changed with day length (Fig.8g). To project these results for any given day length after the summer solstice, the chances of temperatures dropping below zero was almost always smaller for Coldfoot than Toolik, with Sagwon staying in between – there was up to a 28 percentage point difference between Coldfoot and Toolik, which happened when day length dropped to ca. 13 h (Fig.8g). The most striking difference across sites emerged when we computed the number of times freezes occurred when the sun was at or above the horizon (day) versus below the horizon (night). The southern almost doubled the northern sites in night time freeze events, with 67% of freezes occurring at night in Coldfoot, while 37-38% in Toolik and Sagwon respectively (Fig.9, top). Lastly, the decay in cumulative positive degree hours (CPDH) following the summer solstice differed greatly between Coldfoot and the two northern sites (Fig.8i). For example, after the sun started dipping below the horizon the proportion of the actual growing season that remained increased from 45% in both northern sites to 51% in Coldfoot (Fig.8i). Moreover, the CPDH during the period between each freeze event and the end of the PGS in Coldfoot bore no relation to whether the freeze occurred at night or during the day (Fig.9, bottom). Conversely, in Toolik, and even more so in Sagwon, the amount of CPDH that followed night freezes was only a small fraction of that which followed freezes that take place during the day (Fig.9).

**Figure 9.**
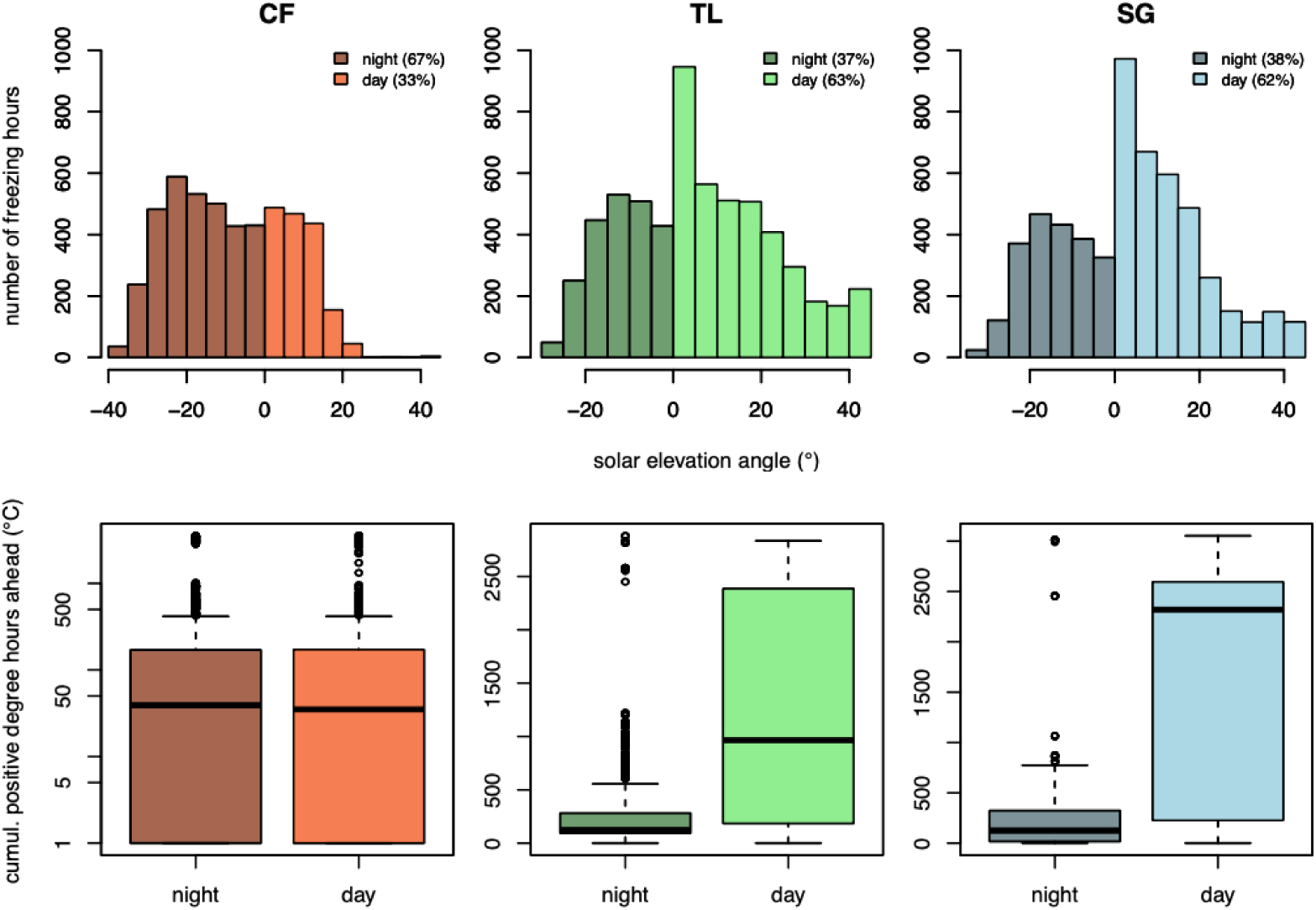
Co-variation between freeze and light regimes in each study site. Top panels show the number of freezing hours occurring at different solar elevation angles during the potential growing season. The proportion of freezing hours occurring while the sun is at or above the horizon (day) as opposed to below the horizon (night) is also shown. Bottom panels show the cumulative positive degree hours (sum of hourly temperatures above 0°C) ahead of each freezing hour that occurred during either the day or night.

### Best performing method to detect the onset of leaf senescence

Among the five algorithms we trained for detecting the onset of leaf senescence (OLS), *maximum drop threshold* was the one that best correlated with the manually picked values of OLS. This algorithm outperformed the others by a large margin (Fig.S1), giving the most accurate and consistent OLS estimates, with R^2^ = 0.45 and root mean squared errors of 8.44. For optimized parameters and detailed performances of all algorithms, see table S3. The *first derivative* and *regression* algorithms yielded moderate performances, while the *second derivative* showed higher error, but the *cumulative sum* approach performed the worst, with large deviations from observations (Fig.S1). Not only were OLS estimates derived by the *maximum threshold* method very close to the manually picked values, but the distribution of absolute errors was narrow, with most predictions falling within ±10 days of observed values (Fig.10). This error range is very similar to the frequency that plants were monitored in the field (ca. 8 days), meaning that the uncertainty in the algorithm estimates is comparable to the uncertainty in the observations themselves.

**Figure 10.**
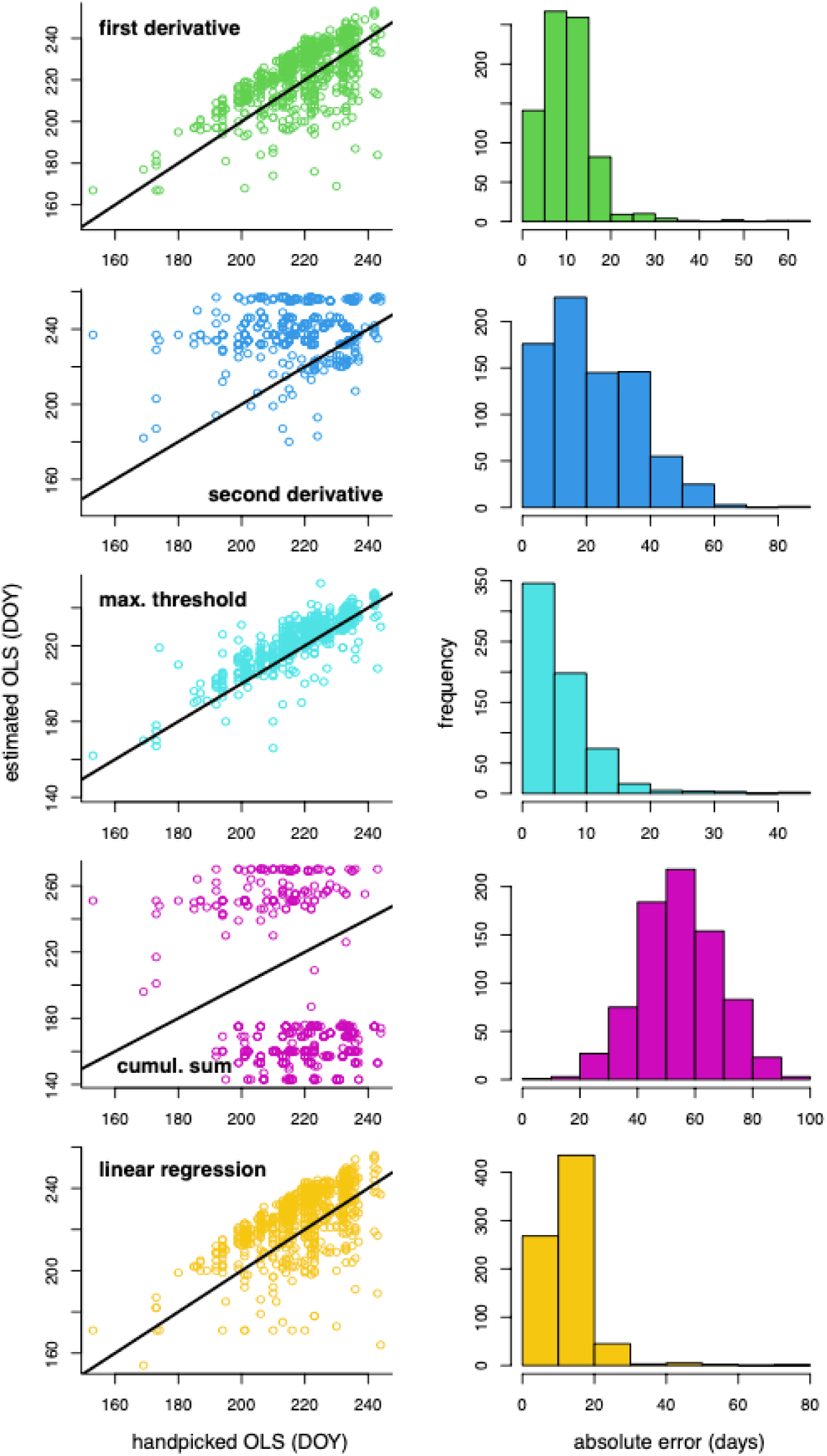
Performance of each of the five algorithms used to detect the onset of leaf senescence (OLS) for each monitored tiller in *Eriophorum vaginatum* plants. The left panels contrast handpicked OLS and the OLS estimated by each algorithm. The black diagonal line is a 1:1 curve. The right panels show the distribution of absolute errors in days.

### Onset of leaf senescence across gardens and ecotypes

We found striking patterns on the dates plants started to senesce (OLS) at the end of the growing season that reflected previous studies (Parker *et al*., 2017; Parker *et al*., 2021). Two patterns are readily distinguishable: 1) northern ecotypes senesce earlier than the southern one in all of the common gardens; and 2) all ecotypes tended to senesce later in the northern gardens than in the southern one (Fig.11, top). These differences in OLS were confirmed by the “group means” hierarchical Bayesian model (Fig.11, bottom). There was also an interaction between ecotype and garden, as the differences among ecotypes varied for each garden, particularly in Sagwon where there were no significant differences among ecotypes (Fig.11). Although our “effects” model did not fit the data as well as the “group means” model (Tab.2), it confirmed this interaction effect (Fig.12). The “effects” model also confirmed that: 1) plants from higher latitudes tend to senesce earlier (5 days per latitude degree), independently of where they were transplanted; and 2) plants at higher latitudes tend to senesce later (4.3 days delay per latitude degree) regardless of their site of origin (Fig.12), which was not specifically measured in previous studies. Lastly, the partitioning of variance in OLS estimated by the “group means” model showed that variation in OLS between years for each sampled tiller was larger than the variance explained by all the other factors combined (Fig.13). We also found a small inter-tiller and inter-plant variance in OLS, compared to the variance explained by ecotype and garden, indicating that plants coming from the same site had very similar responses (Fig.13).

**Figure 11.**
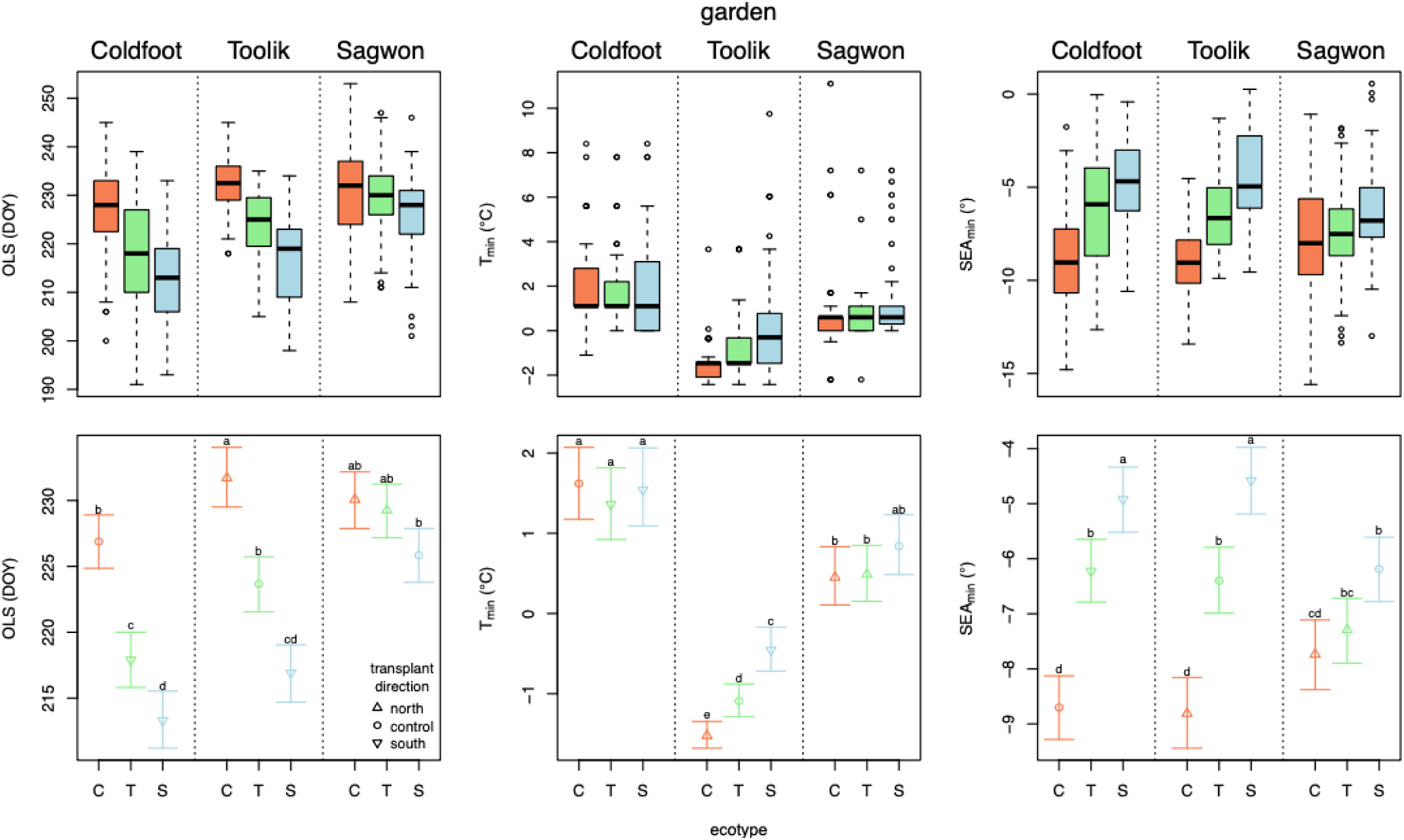
Day of the year (DOY) at the onset of leaf senescence (OLS) of the *Eriophorum vaginatum* individuals used in the reciprocal transplant experiment. Also shown is the minimum air temperature (T_min_) and minimum solar elevation angle (SEA_min_) during the time of OLS. The top panels show the raw data used in the analysis and the bottom panels show means and 95% credible intervals estimated by the “group means” hierarchical Bayesian model. The source population of the plants (ecotype) is denoted by the initial letter of each site (Coldfoot, Toolik, and Sagwon). Compact letter display is used to show groups that are significantly different from one another, i.e., the probability of the range of estimated values overlapping is lower than 5%. Point shape indicates whether plants stayed (control) or were transplanted to the south or north of their sites of origin as shown in the legend.

**Table 2.** Explanatory power (R^2^) of the “effects” and “means” models for each response variable: the onset of leaf senescence (OLS), minimum temperature at OLS (T_min_), and minimum solar elevation angle at OLS (SEA_min_). R^2^ here was estimated based on predictions made for each individual plant and thus ignores variation across years and among tillers in a same plant. CI_95%_ is the 95% credible interval for R^2^.

|  | **Response** | **R<sup>2</sup>** | **CI<sub>95%</sub>** |
|---|---|---|---|
| *Effects model* | OLS | 0.49 | [0.47,0.50] |
|  | T<sub>min</sub> | 0.15 | [0.11,0.17] |
|  | SEA<sub>min</sub> | 0.36 | [0.34,0.38] |
| *Means model* | OLS | 0.59 | [0.57,0.60] |
|  | T<sub>min</sub> | 0.47 | [0.44,0.50] |
|  | SEA<sub>min</sub> | 0.48 | [0.45,0.50] |

**Figure 12.**
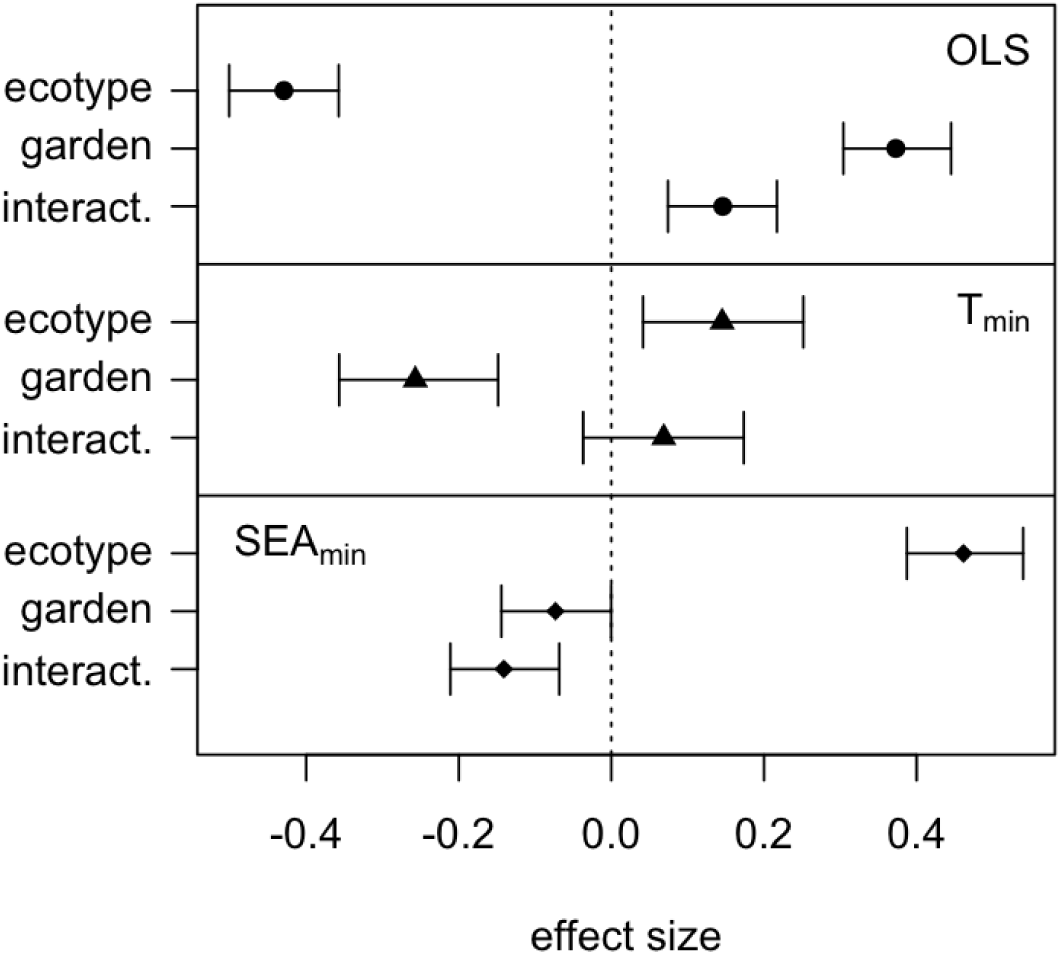
Effect sizes of source population (ecotype), common garden, and the interaction between these two factors on the onset of leaf senescence (OLS), as well as the minimum temperature (T_min_) and minimum solar elevation angle (SEA_min_) at the OLS. Error bars represent the 95% credible interval and points are the median of the posterior distribution for each parameter according to the effects size model.

**Figure 13.**
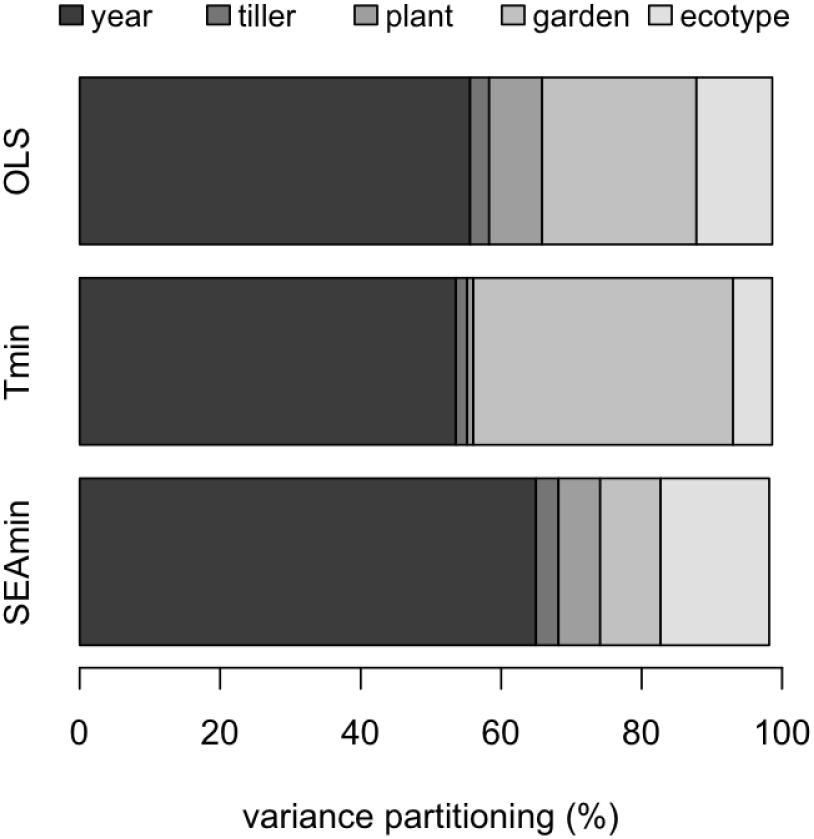
Partitioning of variance in the response variables for the group means model. In this hierarchical model the onset of leaf senescence (OLS), as well as the minimum temperature (T_min_) and minimum solar elevation angle (SEA_min_) at the OLS, were modeled as varying across years (2022-2024), tillers, individual plants, common gardens, and source populations. The credible intervals of each of these variances are shown in Fig.S2.

### Environmental context at the onset of senescence

When we looked at the environmental conditions at the onset of leaf senesce (OLS) we found that minimum air temperature (T_min_) underlay differences in OLS across sites, while within-site differences among ecotypes were better explained by minimum solar elevation angle (SEA_min_). These links are seen in both the “group means” and the “effects” model parameter estimates (Fig.11-13). T_min_ at the OLS was highest in our warmest site (Coldfoot) and lowest in our coldest site (Toolik), with few differences across ecotypes within each garden, except for Toolik, where T_min_ still differed significantly among ecotypes (Fig.11). SEA_min_, in turn, varied widely among ecotypes in Coldfoot and Toolik, but not in Sagwon (Fig.11). Interestingly, in the case of SEA_min_ the ranking was similar across sites, with plants from Coldfoot showing a much lower day length threshold, followed by plants from Toolik and then Sagwon. In other words, day length at the OLS followed the latitude of origin of these plants (Fig.11). These patterns were supported by the results of the “effects” model, which found significant effects of the latitude of plants’ destination (common garden) and origin (ecotype) on the OLS (Fig.12). Whereas gardens had the most effect on T_min_ at the OLS, the ecotype effect was the most pronounced for SEA_min_ (Fig.12). Once again, these patterns could be seen on the partitioning of variance as estimated by the “group means” model, with garden explaining most of the variance in T_min_ and ecotype accounting for most of the variance in SEA_min_, after accounting for inter-year variation in both cases (Fig.13).

When we compared the OLS with the EOS, we found that in Coldfoot control plants start to senesce much earlier (>2 months) than the EOS compared to the northern sites (Fig.S3). This pattern was further confirmed when we calculated the cumulative positive degree hours left between the OLS and the EOS (Fig.14a). On the other hand, control plants in Coldfoot and Sagwon were much less exposed to freezes than in Toolik (Fig.14b). The ecotype specific behavior was most notable in two cases: 1) the Coldfoot plants transplanted in Toolik started to senesce later in the season than at their homesite and were thus exposed to the freezing risks not encountered before transplant; and 2) plants from Sagwon and Toolik transplanted into Coldfoot started to senesce much earlier than the local plants, thus reducing their PGS compared to their homesite (Fig.14c). Another important insight from crossing temperature and OLS data is that Sagwon is the most benign of our sites, with comparatively low freeze risks during the PGS, while Toolik had the highest freeze risk during the PGS (Fig.14c). In summary, these results clearly show that environmental conditions (temperature and day length) are clearly correlated with the OLS of ecotypes across the common gardens.

## Discussion

Given the urgent need to understand how arctic tundra plants will respond to – and whether they can adapt to – climate change, we tested how a foundation arctic species reacted to seasonal changes in temperature and light regimes. In monitoring the late-season leaf phenology of *E. vaginatum* in three reciprocal transplant gardens, we hypothesized that established ecotypes from the taiga and tundra leaf senescence was triggered by either temperature or day length thresholds following experimental results performed 7-9 years prior that did not explicitly address this issue (Parker *et al*., 2021). We found that the three sites where the common gardens were established were each unique in how their freeze regimes were related to seasonal changes in day length. Comparative to the tundra site, the taiga site (CF) had frequency of freezes become common weeks after nights started experiencing full darkness. The southernmost, but highest elevation tundra site, had the most freeze-prone summer, and the percentage of freeze events increased as soon as day lengths became shorter than 24 h. This indicates that, if *E. vaginatum* uses day length as a cue (H_2_), plants coming from southern populations would actually have lower day length thresholds, i.e., be adapted to require more hours of darkness as a senescence cue compared to their northern counterparts. Given that nights reach full darkness earlier in the south, one could predict that: 1) plants would start senescing later when they are moved north, and would senesce earlier when moved south; and 2) in each common garden northern ecotypes would senesce earlier than southern ones. These hypotheses were largely supported by the results of our reciprocal transplant experiment and previous studies in the area (Parker *et al*., 2017; Parker *et al*., 2021). Furthermore, we found that while the temperature at the onset of leaf senescence (OLS) varied the most across common gardens, the day length at the OLS varied the most across ecotypes (within gardens), which has not previously been examined. Therefore, these results shed light on the mechanisms driving plants’ responses to seasonal environmental changes, and thus permit a better understanding of how local adaptation has evolved in these plants.

### The selective pressures along an environmental gradient

The analysis of the freeze regimes across the three sites provided insight into how selective pressures on plant phenology may vary over time and space. Despite occasional mid-summer freezes, the shoulder seasons (spring and fall) clearly constitute the coldest times for Arctic plants during the PGS. This was true across all sites, although spring tended to be more benign in the southern site (Coldfoot), where soon after spring snow melt there were relatively few freeze events. Meanwhile, for the two northern sites (Toolik and Sagwon) the risk of freezes continues to be non-negligible until the summer solstice. At the same time, there are no significant day length cues in the northern sites that plants can use during the spring because snow tends to melt after the 24-h daylight period has already started. If a light-cued mechanism were to evolve, it would be much more likely to happen for leaf senescence, and not for sprouting, which would require other cues like snow melting or an increase in air temperature (Collins *et al*., 2021). The problem with temperature cues to end seasonal growth is that weather is naturally quite variable, both during the course of a single growing season and across years – an issue that tends to be aggravated by the worsening of extreme weather events (Zhou, Leung and Lu, 2024). Finally, the selection for phenological responses to avoid frost damage is likely to be weaker in the spring, when plants have little active tissue to be damaged compared to when they reach peak greenness later in the season.

Because temperature regimes can be unpredictable and the cost of frost-damage increases towards the fall, tundra plants could adapt more reliable environmental cues such as day length to inform their late-season phenology (Fracheboud *et al*., 2009; Gill et al., 2015; Bernhardt *et al*., 2020). However, temperature can have a complementary role, even when photoperiod is a primary driver of senescence in a species (Fracheboud *et al*., 2009; Lang *et al*., 2019). In *E. vaginatum* potential for minimum temperature can theoretically be used by plants to predict near-future freezes: by the time night temperature drops to 5°C this risk is already at ca. 10% (Fig.8f). Minimum daily temperature also indicates how much PGS still remains (Fig.8h). The relationships between temperature and freeze risk for PGS is almost the same across the three studied sites. This lack of spatial variability would not create the selective drive for local adaptation to take place if plants relied only on temperature signals to guide their phenology. On the other hand, light regimes are tied to temperature regimes in a very site-specific manner, and seasonal variation in day length can indeed inform plants of both the risk of freeze events (Fig.8g) and the proportion of the remaining growing season (Fig.8i). This spatial variability could create conditions for the evolution of ecotypes with unique responses to light cues, even though such cues only remain adaptive if the climate does not change (Bernhardt *et al*., 2020) or the ecotypes do not migrate north or south (Ettinger *et al*., 2021). For example, if a northern ecotype migrates south it will still avoid frost-damage very efficiently if it senesces soon after nights have first full darkness, but would be missing a great deal of its PGS. This may in part explain why Bennington *et al*. (2012) found that *E. vaginatum* transplants from the tundra in these same gardens had increased death rates after 30 years when moved to taiga gardens in the south. Likewise, although a southern ecotype might be able to extend its PGS by senescing only after longer periods of darkness in the north, it could risk frost-damage. However, the same long-term studies by Bennington *et al*. (2012) found that southern ecotypes moved into tundra gardens sometimes outperformed or had similar survival as transplanted northern plants in these same gardens, suggesting at least regarding long-term survival, the exposure to these lower temperatures before senescence may not be as detrimental.

The site-specific coupling of temperature and light regimes likely causes local populations to evolve unique phenological responses to individual environmental cues or even combinations of cues. The greatest qualitative environmental difference across sites comes from crossing light and temperature data: when freezes occur during dark nights in the north there is very little of the potential growing season left, while in the south freezing, dark nights do not provide a direct link to the time plants still have left to grow (Fig.9). Thus, there could be selection for an integrated response to both changes in light and temperature in the northern ecotypes, but lacking or more based on cumulative effects of freezing or dark hours in the southern ones. Such integration of distinct cues is consistent with the dual role of phytochromes as both thermal sensors and photoreceptors (Li, Liang and Liu, 2022). Overall, the environmental data suggest that plants face site-specific selection pressure to integrate temperature and/or light cues as triggers of their phenological behavior.

### Using machine learning algorithms to detect the onset of leaf senescence

Out of the five algorithms used to detect the onset of leaf senescence (OLS), the *maximum drop threshold* method performed the best, with an error of just 8.4 days, which indicates that most of the predictions fall within the observational period of measuring green length (7-10 days). Predictions from this model can thus be confidently used in place of manually determining the OLS from observational data. The other algorithms performed more poorly when compared to the *maximum drop threshold* method, likely because they tend to amplify noise more than the *maximum drop threshold*. If there is a plateau in total green length after senescence or even if the decrease in green length slows down but is still negative, the derivative and regression methods will overestimate the importance of these features and remove them from the set of days that could be the start of senescence. *Maximum drop threshold*, on the other hand, always looks at points after the peak green length, and only requires a sustained decline rather than a consistent specific slope. This makes the method more robust to variation in the pattern of decline and better aligned with the biological process of senescence. Change-detection methods that rely on the largest derivative or curvature inflection points are highly susceptible to local fluctuations and noise because the fitted curves are continuous in the first derivative but still carry residual variability from smoothing, such as from sampling frequency, leaf die-off, or variability in measurements (Zeng *et al*., 2020).

A commonly used approach to detecting the OLS is to fit a double logistic curve to the entire seasonal growth trajectory and then select an inflection point as the OLS (Parker *et al*., 2021). While this method is designed to capture the overall shape of the green-up and -down curve during the growing season, it often requires eliminating a number of poorly fitting curves, which reduces its applicability. Compared to this, the *maximum drop threshold* method is more direct and robust, as it is specifically designed to capture a single turning point on the greenness curve. It does not depend on a global parametric fit and instead focuses only on the post-peak decline, making it more usable for noisy seasonal growth data. End of the season senescence transitions are inherently less distinct and more prone to noise than spring green-up, making inflection-based logistic approaches on a global parametric fit less reliable under noisy decline patterns (Zhang *et al*., 2006).

### Adaptation to local climates underlined by phenological responses

Our study showed that plants’ late season phenology depends on both their population of origin and the site where they were planted, indicating *E. vaginatum* populations have adaptations to the local climates where they occur. These results confirm earlier studies coming from the same garden (Parker *et al*., 2021) or done in the same area (Shaver, Fetcher and Chapin, 1986; Fetcher and Shaver, 1990; Bennington *et al*., 2012; Parker *et al*., 2017) showing that *E. vaginatum* forms classic ecotypes along its distribution range, i.e., local populations that can be clearly distinguished by the unique way they respond to the environment (Stebbins, 1950). This study also contributes to the long-standing tradition of common garden and reciprocal transplant experiments in plant evolution, even though only a minority of such studies focuses on plant phenology (Johnson *et al*., 2021). This scarcity of phenological monitoring on reciprocal transplants is intriguing given that one of the most striking results of the original experiment was the distinct phenological behavior shown by different ecotypes of *Drymocallis lactea* (Clausen, Keck and Hiesey, 1940). This study thus fills an important knowledge gap in the plant evolution literature, a gap that is becoming increasingly important as climate change threatens the very existence of local populations of many widespread species (Peterson, Loak and Morris, 2018). To that effect, this study provides an apparently paradoxical lesson. Taken at face value, it displays the limitations in the ability of reciprocal transplants to predict the effects of climate change, as the unique combination of light and temperature regimes at each site prevents generalizations across the species distribution range. At the same time, our study demonstrates the potential for these experiments to shed light into the mechanisms driving local adaptation, which in turn are generalizable and may be used to predict climate change impacts on local populations.

In our experiment, having plants from contrasting habitats grown together under the same environmental conditions allowed us to test for local adaptation. We found that plants coming from different populations can show quite contrasting phenological behaviors even after 8-10 years growing in common gardens. This directly contradicts the possibility that all ecotypes could be pre-adapted to respond to a single temperature threshold to cope with different local climates (H_0_ in the conceptual framework section). Another implication is that these source populations have genetic (Johnson *et al*., 2021; Stunz *et al*., 2022) or epigenetic (Robertson and Wolf, 2012) adaptations to local climates. This conclusion aligns with landscape genomics evidence that points to genetic structure dividing populations north (Toolik and Sagwon) and south (Coldfoot) of the treeline (Stunz *et al*., 2022), where the strongest difference in end of season phenology is detected. Our results, however, also point to clear differences in phenological behavior between the northern populations, especially when grown in the Toolik garden (Fig.11). These phenological differences may represent the different climate between these two sites, with Toolik being more prone to summer freezes and having a shorter potential growing season than Sagwon (Fig.8). Therefore, if there is a genetically mediated senescence response between these populations it will likely be best determined with finer scale genomic or transcriptomic data. Our results also demonstrate that common garden experiments conducted in isolation can be insufficient and even misleading. For example if only the garden in Sagwon (Fig. 11) was examined, the lack of strong differentiation among the three ecotypes in phenology would have underestimated the importance of ecotype adaptation.

The phenological variation observed across common gardens and ecotypes allowed us to constrain the set of potential mechanisms driving local adaptation in this species. First, the order in which ecotypes started to senesce reflected the latitude of their sites of origin – even in Sagwon, where these differences were not significant, the order was still the same – with the northern ecotypes always senescing before the southern ecotype (Fig.11). This does not support ecotype-specific temperature thresholds (H_1_) for senescence. However, these results do support the hypothesis of a light mediated response (H_2_) for senescence as has been shown in many lineages of plants (Jiang *et al*., 2022), especially at high latitudes (Gill *et al*., 2015). Moreover, the order at which ecotypes started to senesce in each garden also corroborated the idea that northern ecotypes respond to longer day length thresholds than southern ecotypes (H_2_). There has indeed been some experimental evidence showing *E. vaginatum* growth and fecundity to be sensitive to seasonal changes in day length (Shaver and Kummerow 1992). However, this might be the first explicitly tested evidence that changes in day length cue leaf senescence of this important arctic species. In spite of the overwhelming support for day length being the major factor driving the onset of leaf senescence in *E. vaginatum*, there are still some results that cannot be explained by the effects of day length alone.

We found indirect evidence that temperature thresholds may play a supporting role in triggering leaf senescence. Our best supported hypothesis (H_2_) could not explain the convergence of the onset of leaf senescence of all three ecotypes towards the northernmost site (Fig.11). This convergence clearly indicates an interaction between ecotype and common garden effects, which was supported by our effects model (Fig.12). This interaction would be expected if phenological responses to temperature were modulated by day length – plants would become increasingly sensitive to low temperatures as day length shrank towards the end of the growing season (H_3_). H_3_ also predicts that there should be a reversal in the order ecotypes senesced; this might have not happened only because the latitudinal range we used in our study did not establish gardens far enough north. The interactive effect of light and temperature on plants is actually expected given that the main class of integrator molecules responsible for controlling leaf senescence (phytochromes) are both light receptors and thermosensors (Li, Liang and Liu, 2022) and models for understanding the cumulative effects of day length and temperature are being developed (Lang *et al*., 2019). An example of how light and temperature may interact to elicit phenological responses is found in *Picea abies*, whose bud burst is only promoted by a combination of rising temperatures and lengthening day lengths (Partanen, Koski and Hänninen, 1998). In our study, a more prosaic explanation for the convergence of leaf senescence onset at our northernmost site is that many plants, primarily from the southern ecotype, remained green by the time phenology monitoring had to be terminated at the end of the growing season.

Nevertheless, day length has such a dominant effect over temperature in triggering leaf senescence in *E. vaginatum* that, even though our middle site (Toolik) is actually colder than our northernmost site (Sagwon), the order of senescence followed instead the latitude of the sites of origin. These common garden results support the hypothesis of sensitivity to hours of darkness mediating senescence response. This mismatch caused Toolik plants to have the highest exposure to freezes when in their home habitats compared to the other two control groups (Fig.14). This result seems incompatible with the assumption that ecotypes are optimally adapted to their local climates, so we entertain two post-hoc hypotheses. The first is that in recent decades Toolik has not warmed as fast as Coldfoot and Sagwon, thus creating the impression that the Toolik ecotype is more frost-tolerant than the others. This is on par with the notion that higher elevation sites might constitute climatic refugia as it has been shown in the boreal biome (Stralberg *et al*., 2020). Alternatively, the Toolik ecotype might actually have evolved to become more freeze-resistant in order to better exploit the much shorter growing season compared to the other sites. It has indeed been shown for multiple species in the Alps that ecotypes from higher elevations are considerably more freeze-resistant than their lower-elevation counterparts (Neuner *et al*., 2020). Independently of what explanation is correct, it is clear that the eco-physiological context in which local adaptation takes place is more complex than anticipated.

**Figure 14.**
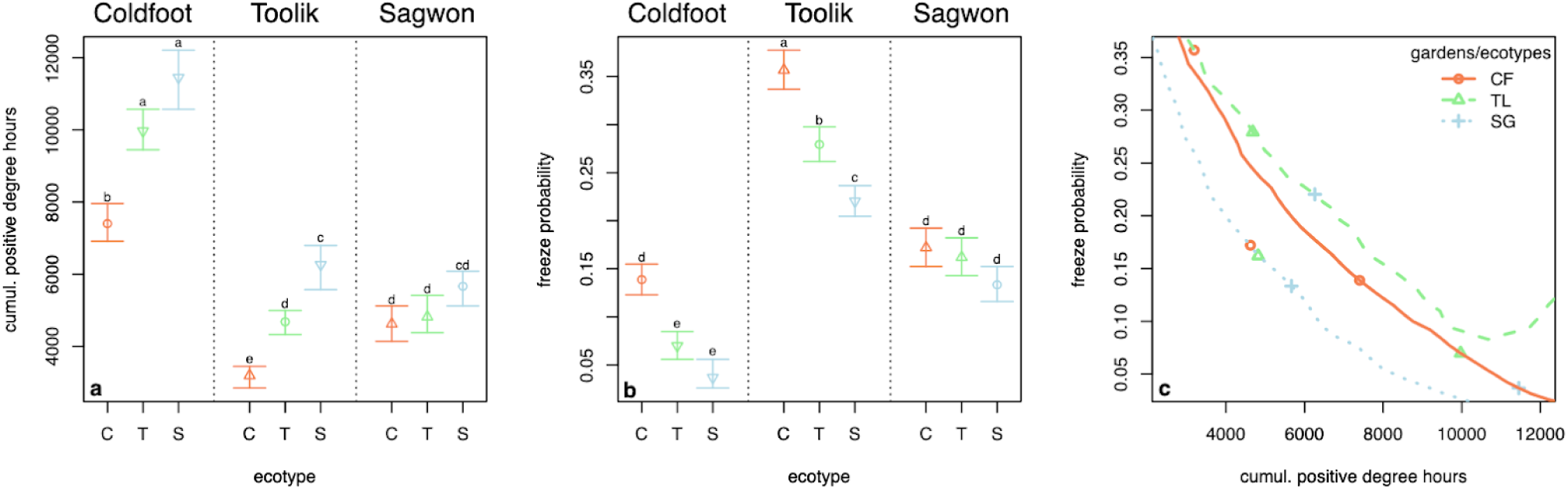
Plant ability to exploit the potential growing season and the freeze exposure cost that comes with it. Cumulative positive degree hours (CPDH) left between the time plants start to senesce (OLS) and the end of the potential growing season (a) and the probability of air temperature falling to or below freezing point (b) across sites and ecotypes. Crossing this information yields a space representing the trade-off between conservative (low freeze exposure) and acquisitive (ability to exploit the season) phenological strategies (c). These graphics are based on the OLS estimated by the “group means” model, and freeze probability and CPDH are averaged over the 2012-2024 period. CPDH and freeze probabilities at the observed OLS are shown in Fig.S4.

Local adaptations for senescence in this system appears to be based on the phenology-mediated trade-off between avoiding freezes and exploiting the potential growing season. The timing of leaf senescence could be seen as a one-trait trade-off, in which the trait value being selected varies across habitats precisely due to opposing selection forces (Agrawal, Conner and Rasmann, 2010) – minimizing frost-damage and maximizing competitive ability. That means that for each site there comes a time towards the end of the summer when the cost of frost-damage starts to surpass any growth gains from overextending the growing season. Whatever day length happens to be in place at this critical time may be used by plants as a cue for senescence. Therefore, given that warmer, southern sites will get dark earlier and remain warmer longer than those in the north, plants in the south that remain active for a longer time after it starts to get dark at night would be favored. Meanwhile, colder, northern sites should favor plants whose senescence is triggered by less hours of darkness before the environment becomes unsuitable. This is evidenced in these transplant gardens where the plants retain phenological characteristics akin to their homesites, but that may no longer be adaptive: while the southern ecotype transplanted north may get much more exposed to freezes than local plants, northern ecotypes transplanted south start senescing prematurely compared to the local ecotype (Fig.14c). In the context of this study, this one-trait trade-off can be seen as a spatial niche partitioning mechanism (Amarasekare, 2003), with more dark-sensitive ecotypes being restricted to higher latitudes and less dark-sensitive ecotypes restricted to lower latitudes within *E. vaginatum*’ distribution.

Within sites, however, this selective process is believed to be stabilizing (Agrawal, Conner and Rasmann, 2010), which translates into reduced local variability. Indeed, variation in the onset of leaf senescence among individuals of the same ecotype was only not smaller than the variation among the tillers of an individual plant (Fig.13). This lack of local variation in phenological strategies might, in turn, compound the risk posed by a rapidly warming Arctic. Our results suggest that with climate change *E. vaginatum* will continue to senesce prematurely even as the growing season continues to extend later. On the other hand, if southern ecotypes could move north they would have the potential to be better adapted to the changing tundra climate. However, there is a gene flow resistance barrier at the treeline that could limit movement in combination with the presence of long-lived northern ecotypes that occupy the tundra (Stunz *et al*., 2022) that would make replacement a significant challenge.

### Next steps: further analyses, data, and methods

Even though the findings in this paper are quite striking, there is still much to learn about this system, so we here offer some potential next steps. These steps can be taken by exploring new analytic approaches, collecting or compiling new data, and designing new experiments. For instance, the database used in this study – weather and leaf phenology data – is too cumbersome to be tackled at once, so many simplifying assumptions had to be made for the sake of tractability. Perhaps the main one was to reduce our phenology data into a single metric (onset of leaf senescence; OLS): not only there is a lot to be learned about the green-up and post-OLS phases, but leaf-level data could generate further insights into the factors that drive leaf expansion and senescence. Another topic to be further explored that was evident from our results is the wide year-to-year variation in the phenology of individual tillers, which could show how climatic variability and change can further affect this species’ phenology. In short, a more detailed analysis, done at the leaf-level, for the entire phenological trajectory, and using climatic data as predictor variables should be tried, probably by employing machine learning techniques given the complexity of the analysis.

This prolific study system could also greatly benefit from some key, new data on local climates and on the plants themselves. For instance, in this study we assumed day length was the main way light could affect plants’ phenology, but perhaps light intensity also plays a role and could explain part of the great year-to-year variation in phenology. Prolonged cloudiness towards the end of the growing season could mimic the effects of shortening day length, thus misleading plants into senescing earlier. As for the plants, there are some exciting new avenues that can be explored: 1) whole-plant fitness metrics could further test our conclusions about local adaptation and the misfit of transplanted individuals; 2) the use of phenocams could grant much improved temporal resolution (daily phenological data) and a much wider spatial extent, as they do not require a team to be frequently visiting each study site; and 3) genetic studies could elucidate the physiological pathways of leaf senescence in this particular species. The use of phenocams in particular would allow us to more properly test our hypotheses by greatly expanding on the climatic range being sampled, and by including ecotypes with even more contrasting genetics. Further genetic studies focused on differential gene expression among ecotypes before and after the onset of leaf senescence (transcriptomics, e.g.) could shed some light into the molecular mechanisms driving plant phenology. By narrowing down on the myriad of potential physiological pathways plants can take to regulate their phenology we could better understand the evolutionary processes driving local adaptation. Lastly, factorial experiments manipulating photoperiod and temperature regimes would allow us to properly test the causality we can only infer from field studies like this. Although field experiments would already be a great advance, it is not possible to fully disentangle light from temperature in the field, as they are usually too correlated. Only lab experiments conducted in controlled settings could allow us to test the individual and combined effects of light and temperature regimes on the onset of leaf senescence.

## Conclusion

Coupling between freeze and day length regimes can be a crucial selective force driving the evolution of local adaptation in *Eriophorum vaginatum*. The primary mechanism by which populations adapt to local climates appears to be the triggering of leaf senescence by site-specific day length thresholds, although temperature cues seem to play a minor role in these responses. This study provides an important step towards more accurate predictions of the immediate effects climate change will have on arctic tundra vegetation. As summer temperatures rise and new patterns of freezing emerge, the reliance of plants on predictable cues such as day length may become maladaptive. If light and temperature regimes become decoupled, local populations may not be able to exploit expanding growing seasons and arctic foundation species may lose ground to more competitive species, such as local shrubs and plants migrating from lower latitudes. If that is the case for *E. vaginatum*, as this study indicates, assisted migration and/or assisted gene flow from southern ecotypes, despite the risks involved (Aitkin and Whitlock, 2013), might become a legitimate option to preserving the vasts areas covered by tussock tundra in Northern Alaska and the huge carbon stocks they hold (Curasi *et al*., 2022).

## Acknowledgements

We are deeply grateful for all the support this work has received from the staff from the Toolik Lake Field Station, especially from the Environmental Data Center team, who compiled weather data supported by NSF grants OPP-2109975, OPP-2109946, and OPP-2109950. We also thank Christina Rinas for help with setting up the reciprocal transplant experiment in 2014 as well as the undergraduate students who helped us in the field between 2022 and 2024 (Allie Faunce and Lawrence Tang). We acknowledge these data were collected and this research conducted on the unceded lands of the Dena’ina and lower Tanana Dene Peoples in our northern sites, and of the Nunamiut People in our southern site.

## Author contribution statement

Conceptualization: N.F., J.T., M.L.M, G.S., M.C.V.

Data curation: M.C.V., B.L., W.M., H.O. Formal analysis: M.C.V., W.M.

Funding acquisition: M.L.M., N.F., J.T., G.S.

Investigation: B.L., M.C.V., B.D.C.R., L.A.P.S., W.M., C.E., H.O., E.K., D.J., N.F.

Methodology: G.S., N.F., M.C.V., W.M., L.A.P.S., B.L.

Project administration: M.C.V., N.F., M.L.M., G.S., B.L., L.A.P.S.

Resources: N.F., M.L.M., J.T., G.S.

Software: W.M., M.C.V. Supervision: M.C.V. Validation: W.M., M.C.V.

Visualization: M.C.V., W.M., C.E., B.D.C.R. Writing – original draft: M.C.V., W.M., D.J.

Writing – review & editing: M.C.V., W.M., N.F., M.L.M., B.L., G.S., D.J., B.D.C.R., E.K., H.O., C.E., L.A.P.S.

## Data availability statement

For full replicability of this paper, we provide full access to the data and the code used in the analyzes. The compiled climate data, the leaf phenology data, and the R and Python code will all be deposited at the Arctic Data Center (https://arcticdata.io). The OLS detection algorithms, data, predictions, and parameters are also available at https://github.com/milkman1017/Senescence_Detection

## Field research permits

This research was conducted under Alaska Department of Natural Resources permit LAS29722 and Bureau of Land Management permit F-97293.

