## Appendix for "Ecotypes matter: photoperiod and site of origin underlie the onset of leaf senescence in *Eriophorum vaginatum*, an Arctic foundation species"

**Machine Learning Algorithms for Detecting Onset of Leaf Senesecence**

Starting from the smooth leaf data described in the Methods section, here is a more detailed descriptions of the other four algorithms:

1. First Derivative: First we estimate the change in green length over every day:

$$d_t = g_t - g_{t-1} \text{ (Eq. 1)}$$

To try and minimize noise, we apply a rolling average of the green length first derivative values:

$$\bar{d}_t = \frac{1}{N_t} \sum_{i \in W_t^{(d)}} d_i \quad \text{with } N_t \geq p_d \quad \text{(Eq. 2)}$$

Where $W_t^{(d)}$ is the number of days in the rolling window, $N_t$ is the number of valid observations within the window (to handle the window edges), and $p_d$ is the necessary minimum number of points within the window

Next we identify periods of a sustained drop in green length:

$$z_t = \begin{cases} 1 & \text{if } \bar{d}_t < 0 \\ 0 & \text{otherwise} \end{cases} \quad \text{(Eq. 3)}$$

To count as a sustained drop, we required that within a rolling window of days $w_s$, at least $p_s$ valid points were present (again, to handle windows at the edge of the data set), and of these days, at least $\theta_s$ consecutive days showed a decline in green length:

$$y_t = \begin{cases} 1 & \text{if } \sum_{i \in W_t^{(s)}} z_i \geq \theta_s \quad \text{with } |W_t^{(s)}| \geq p_s \\ 0 & \text{otherwise} \end{cases} \quad \text{(Eq. 4)}$$

Finally, OLS is computed as the first day where the drop is sustained for the required minimum number of days, adjusted by some constant offset $\Delta$:

$$\text{OLS}_{f.d.} = \min\{t \mid y_t = 1\} + \Delta \text{ (Eq. 5)}$$

2. Second Derivative: Like the first derivative method, we start by calculating and smoothing the first derivative:

$$d_t = g_t - g_{t-1}$$

$$\bar{d}_t = \frac{1}{N_t} \sum_{i \in W_t^{(1)}} d_i \quad \text{with } N_t \geq p$$ (Eq. 6)

Where $W_t$ is the window size and $p$ is the minimum number of points within the window. Similarly, we find the second derivative on the smoothed first derivative and then smooth with another rolling average:

$$s_t = \bar{d}_t - \bar{d}_{t-1}$$

$$\bar{s}_t = \frac{1}{M_t} \sum_{i \in W_t^{(2)}} s_i \quad \text{with } M_t \geq p$$ (Eq. 7)

$M_t$ is the window size used to smooth the second derivative and $p$ is the number of points within the window. We define the OLS as the first day where the second derivative crosses some threshold value $\tau$, indicating an inflection point

$$t^* = \min\{t \mid \bar{s}_t < -\tau\}$$ (Eq. 8)

The final OLS is adjusted by some constant offset $\Delta$

$$\mathrm{OLS}_{\mathrm{s.d.}} = t^* + \Delta$$ (Eq. 9)

3. Rolling Regression: Here we try a similar philosophy to the first derivative method, but instead of taking the first derivative over just two days, we find the slope over a window of days $W_t$ to hopefully reduce sensitivity to noise. A simple linear regression is fit over a rolling window with at least $p_t$ valid points:

$$g_\tau \approx \beta_0 + \beta_1 \tau, \quad \tau \in W_t$$ (Eq. 10)

And extract the slope $\beta_1$ to obtain a sequence of slopes:

$$s_t = \beta_1(W_t)$$ (Eq. 11)

To reduce noise, slopes that deviate more than some s.t.d. threshold $k\sigma_s$ are or fall below some $R^2$ threshold $\rho$ are removed from the set

$$|s_t - \mu_s| \leq k\sigma_s \text{ (Eq. 12)}$$

$$R^2(W_t) \geq \rho \text{ (Eq. 13)}$$

Then we identify times where the slope is negative and below some value $\gamma$:

$$\mathcal{N} = \{t \mid s_t < \gamma\} \text{ (Eq. 14)}$$

We require at least $L$ consecutive days be below the threshold to be considered senescence:

$$\mathcal{S} = \{t \in \mathcal{N} \mid s_{t'}, \ldots, s_{t+L} < \gamma\} \text{ (Eq. 15)}$$

The start of senescence is the first day in this set:

$$t_{\text{sen}} = \min \mathcal{S} \text{ (Eq. 16)}$$

And the final OLS prediction is adjusted by some constant offset $\Delta$

$$\text{OLS}_{\text{reg}} = t_{\text{sen}} + \Delta \text{ (Eq. 17)}$$

4. Cumulative Sum: From the smoothed green length, we calculate the deviation from the baseline expanding mean at each time point:

$$\bar{g}_t = \frac{1}{w_{\text{base}}} \sum_{\tau=t-w_{\text{base}}+1}^{t} g_\tau$$

$$d_t = g_t - \bar{g}_t \text{ (Eq .18)}$$

And sum their contributions at each point

$$c_t = \sum_{\tau=1}^{t} d_\tau \text{ (Eq. 19)}$$

To reduce noise, we apply a rolling mean of cumulative sum deviations over a window $w_t$:

$$\tilde{c}_t = \frac{1}{w} \sum_{\tau=t-w/2}^{t+w/2} c_\tau \quad \text{(Eq. 20)}$$

And then apply trend smoothing to reduce noise further:

$$\tilde{c}_t^{(\text{trend})} = \frac{1}{w_{\text{trend}}} \sum_{\tau=t-w_{\text{trend}}/2}^{t+w_{\text{trend}}/2} \tilde{c}_\tau \quad \text{(Eq. 21)}$$

Then we define some quantile cutoff $\theta_q$ to determine when this deviation is greater than usual:

$$\theta_q = Q_q(\tilde{c}_t), \quad q \in (0,1) \text{ (Eq. 22)}$$

The total drop in the smoothed cumulative sum change needs to be above some minimum value $\Delta c_{min}$:

$$\max_t \tilde{c}_t - \min_t \tilde{c}_t \geq \Delta c_{\text{min}} \text{ (Eq. 23)}$$

Senescence is determined by the first day where the smoothed cumulative sum deviation trend drops below the quantile threshold, provided the minimum change is also satisfied:

$$t_{\text{sen}} = \min\{t \mid \tilde{c}_t^{(\text{trend})} < \theta_q \ \text{ and } \ \max_t \tilde{c}_t - \min_t \tilde{c}_t \geq \Delta c_{\text{min}}\} \text{ (Eq. 24)}$$

Again, the final OLS adjusted by some constant offset $\Delta$

$$\text{OLS}_{\text{cumsum}} = t_{\text{sen}} + \Delta \text{ (Eq. 25)}$$
