## Supplementary material for "Ecotypes matter: photoperiod and site of origin underlie the onset of leaf senescence in *Eriophorum vaginatum*, an Arctic foundation species": Suppl. material

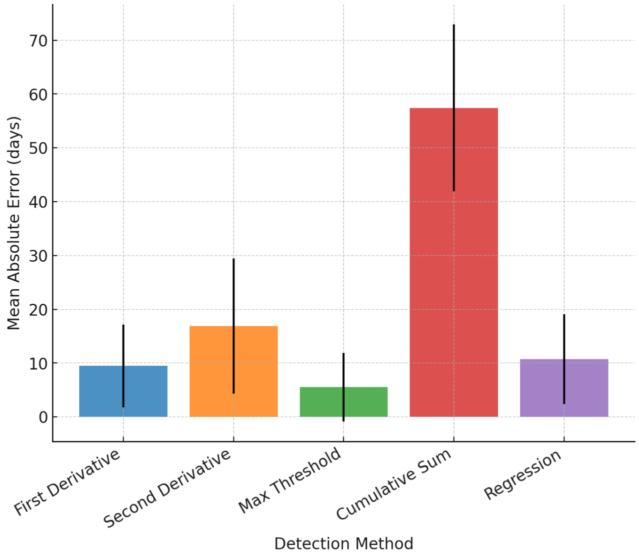

**Figure S1.** Individual Senescence DetectionModel Performance

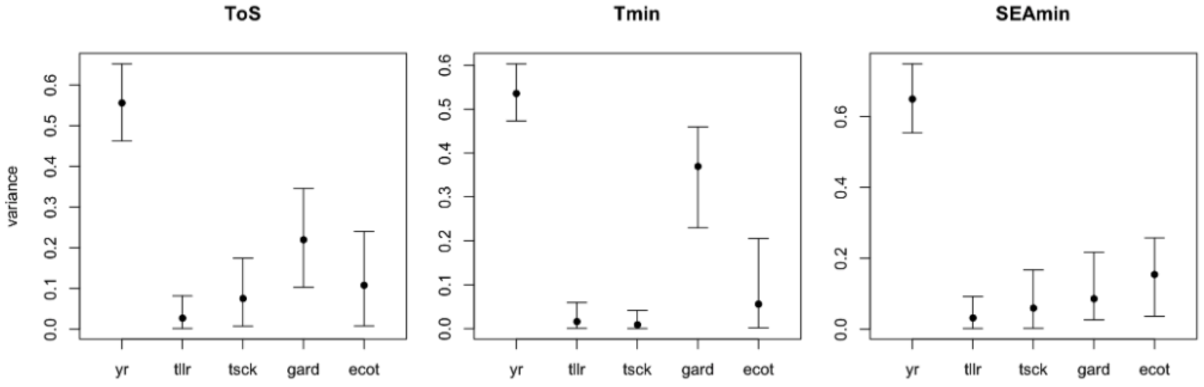

**Figure S2.** Point estimates and credible intervals of the variance parameters used in the group means model.

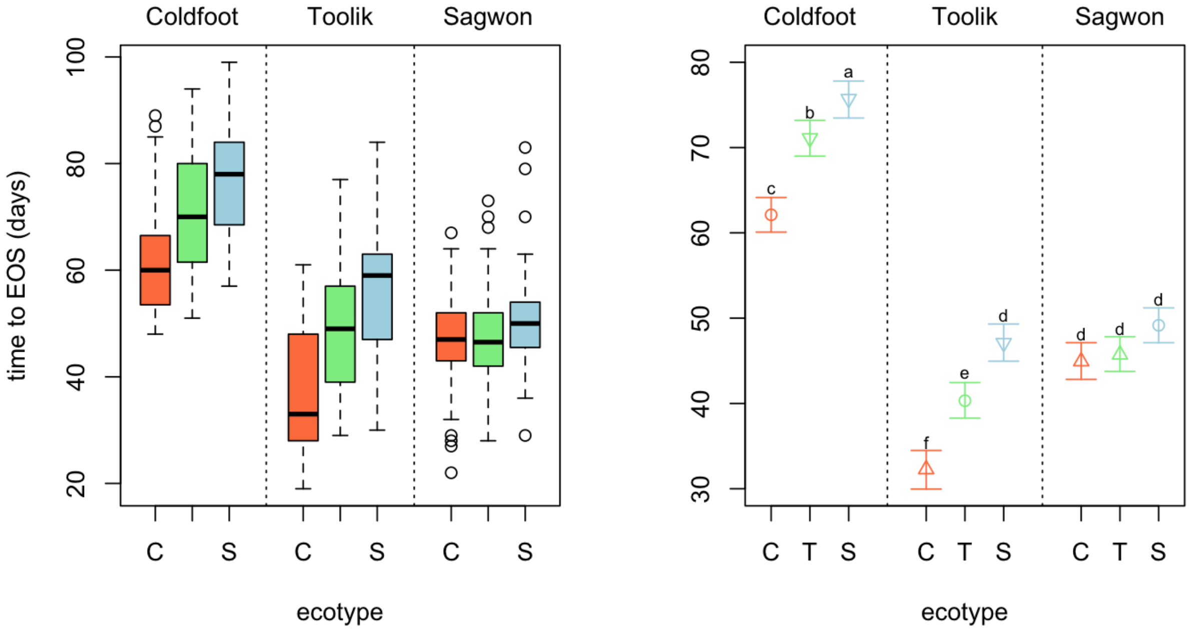

**Figure S3.** Time interval between the onset of senescence, observed (left) and estimated by the "group means" model (right), and the end of the potential growing season in each of the common gardens.

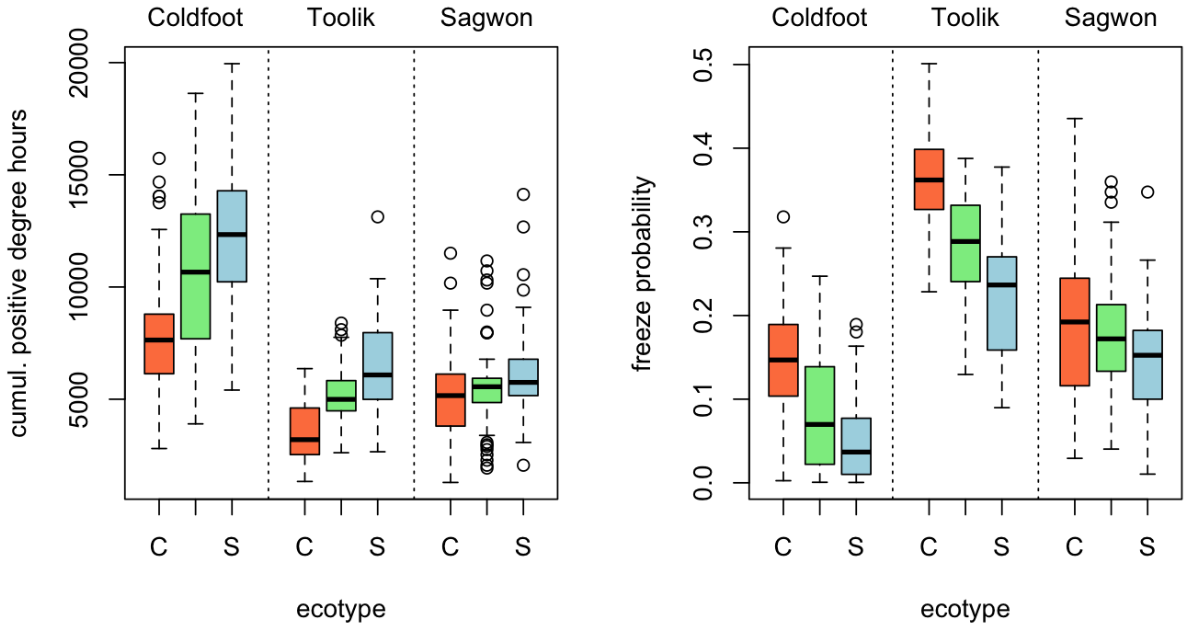

**Figure S4.** Cumulative positive degree hours between the observed onset of senescence and the end of the potential growing season (left). Probability of daily temperatures falling to freezing point or below at the time plants started to senesce (right).

**Table S1.** Monitoring schedule for the monitoring of the vegetative phenology of *Eriophorum* for each year and common garden. Start and end dates are shown as day of the year and the average frequency of visits is given in days.

| Year | Garden | Start | End | Frequency |
|---|---|---|---|---|
| **2022** | Sagwon | 168 | 239 | 8.9 |
| | Toolik | 153 | 241 | 8.8 |
| | Coldfoot | 153 | 244 | 7.6 |
| **2023** | Sagwon | 168 | 263 | 7.9 |
| | Toolik | 164 | 262 | 7.5 |
| | Coldfoot | 146 | 264 | 9.1 |
| **2024** | Sagwon | 162 | 248 | 7.2 |
| | Toolik | 150 | 250 | 7.1 |
| | Coldfoot | 136 | 253 | 7.3 |

**Table S2**. Parameters used and optimized in each of the OLS Machine Learning Algorithms.

| Algorithm | Parameter | Notation | Description |
|---|---|---|---|
| *First Derivative* | First Derivative Window size | $w_d$ | Size of the rolling window (in days) used to compute the first derivative |
| | First Derivative Periods | $p_d$ | Minimum number of data points within the rolling mean of the first derivative |
| | Drop Window | $w_s$ | Size of the window used when detecting sustained drops in the first derivative slope |
| | Drop Periods | $p_s$ | Minimum number of periods needed to detect a sustained drop |
| | Sustained Drop Threshold | $\theta_s$ | Number of consecutive data points where the rolling derivative must be negative to be considered a |

| | | | |
|---|---|---|---|
| | | | sustained drop |
| | Offset | $\Delta$ | Constant value added to the final OLS prediction |
| *Second Derivative* | First derivative window | $W_t$ | Size of the rolling window to compute the first derivative over |
| | Second derivative window | $M_t$ | Size of the rolling window to compute the second derivative over |
| | Minimum periods | $p$ | Minimum number of data points needed to compute the rolling mean for both the first and second derivative windows |
| | Threshold | $\tau$ | Minimum negative value of the smooth second derivative to be considered an inflection point |
| | Offset | $\Delta$ | Constant offset to adjust final OLS |
| *Cumulative Sum* | Window size | $w_t$ | Size of the window used to compute the smoothed cumulative sum |
| | Quantile | $\theta_q$ | Threshold used for identifying a significant drop in the cumulative sum |
| | Trend Window | $w_{trend}$ | Size of the window to detect if the cumulative sum is increasing or decreasing |
| | Minimum cumulative sum change | $\Delta c_{min}$ | Minimum change in the cumulative sum needed to detect senescence |
| | Baseline Window | $w_{base}$ | Size of the window used to compute the baseline |

| | | | |
|---|---|---|---|
| | | | green length that the cumulative sum is compared to |
| | Offset | $\Delta$ | |
| *Linear Regression* | Window size | $W_t$ | Size of the rolling window that linear regression is applied to |
| | Minimum periods | $p_t$ | Minimum number of data points needed to compute the regression within the window size |
| | Sustained Days | $L$ | Number of consecutive days that have a negative slope required to be considered senescence |
| | Minimum Slope | $\gamma$ | Minimum negative slope needed to be considered senescence |
| | $R^2$ threshold | $\rho$ | Minimum r-squared value of the regression fit to be considered valid |
| | Outlier standard deviation | $\sigma_s$ | THe number of standard deviations used to identify and filter out outlier data points within the rolling window |
| | Offset | $\Delta$ | |

**Table S3.** Optimized parameters and performance of each machine learning algorithm.

| **Algorithm** | **RMSE** | **$R^2$** | **Parameter** | **Optimized** |
|---|---|---|---|---|

| | (days) | | | value |
|---|---|---|---|---|
| *Maximum threshold* | 8.44 | 0.45 | Window Size | 5 |
| | | | Threshold Fraction | 0.90 |
| | | | Minimum Drop Duration | 7 |
| | | | Offset | -2 |
| *First Derivative* | 12.22 | -0,17 | Derivative Window | 3 |
| | | | Derivative Periods | 3 |
| | | | Drop Window | 10 |
| | | | Drop Periods | 7 |
| | | | Sustained Drop Threshold | 7 |
| | | | Slope Threshold | -0.05 |
| | | | Offset | 7 |
| *Second Derivative* | 21.07 | -2.34 | First Derivative Window | 7 |
| | | | Second Derivative Window | 7 |
| | | | Min Periods | 5 |
| | | | Threshold | 0.868 |
| | | | Offset | -7 |
| *Regression* | 13.63 | -0.45 | Window Size | 16 |

| | | | | |
|---|---|---|---|---|
| | | | Min Periods | 9 |
| | | | Sustained Days | 9 |
| | | | Slope Threshold | 0.1 |
| | | | R2 Threshold | 0.6 |
| | | | Outlier Std. | 2.5 |
| | | | Offset | 2 |
| *Cumulative Sum* | 59.49 | -25.49 | Window Size | 3 |
| | | | Quantile | 0.01 |
| | | | Trend Window | 21 |
| | | | Minimum Sum Change | 0.2 |
| | | | Baseline Window | 28 |
| | | | Offset | 7 |
